# Chemically encoded pH-tunable covalent adhesion by a bacterial thioester domain

**DOI:** 10.64898/2026.02.05.703941

**Authors:** Yuki Tokunaga, Ryo Matsunaga, Hiroko Kozuka-Hata, Masaaki Oyama, Kouhei Tsumoto

**Affiliations:** Department of Bioengineering, School of Engineering, The University of Tokyo, 7-3-1 Hongo, Bunkyo-ku, Tokyo, 113-8656, Japan; Department of Chemistry and Biotechnology, School of Engineering, The University of Tokyo, 7-3-1 Hongo, Bunkyo-ku, Tokyo, 113-8656, Japan; The Institute of Medical Science, The University of Tokyo, 4-6-1 Shirokanedai, Minato-ku, Tokyo, 108-8639, Japan

## Abstract

Thioester domains (TEDs) represent a family of bacterial adhesin domains that mediate covalent anchoring to target ligands via an intramolecular thioester bond. Although the broad distribution of TEDs among Gram-positive bacteria suggests a critical functional role, the biological significance of this covalent mechanism remains unclear. In this study, we demonstrated that TED-mediated covalent anchoring is reversible and that its equilibrium is regulated by pH. Specifically, SfbI-TED from Group A Streptococcus binds tightly to fibrinogen at physiological pH, whereas mild acidification to pH 6.0 induces rapid dissociation of the complex. Thermodynamic analyses revealed that this pH-responsiveness arises from the intrinsic properties of the thioester bond within the TED. Similar pH-dependent behavior was observed in phylogenetically distinct TEDs, suggesting that pH-responsive adhesion is a conserved feature of the TED family across Gram-positive bacteria.

**Teaser:** A reactive thioester bond in a Gram-positive adhesin acts as a pH sensor to enable environment-responsive adhesion.

## Introduction

Throughout the progression of infection, pathogenic bacteria encounter dynamic changes in the host environment to which they deploy distinct strategies depending on the stage of infection. Among the environmental factors that influence bacterial physiology, pH is a critical determinant of virulence. For instance, Group A *Streptococcus* (GAS) encounters heterogeneous pH microenvironments, typically ranging from mildly acidic (∼pH 5.8) to near-neutral (pH 7.4), across various infection phases, such as initial pharyngeal attachment, biofilm formation, and invasion of deeper tissues (*1–3*). To adapt to these pH variations, GAS utilizes several pH-responsive regulators (*e.g.*, CovR/S (*4*, *5*), CiaH/CiaR (*6*), and RopB (*7*)) to modulate the expression of virulence factors, including adhesins, capsules, streptokinases, and various toxins (*3*, *7*, *8*). Although such gene-regulatory circuits have been the primary focus of research on bacterial pH responses (*7*, *9*), it is equally crucial to elucidate how the function of expressed proteins is modulated by environmental pH. In particular, pH-dependent functional modulation of host-interacting surface proteins, such as adhesins, has been characterized in only a limited number of systems and remains poorly understood (*10–14*).

In this study, we investigated the pH-responsiveness of the thioester domain (TED), a family of adhesin domains prevalent among Gram-positive bacteria across the phyla *Firmicutes* and *Actinobacteria* (*15*, *16*). First reported in 2010 (*17*), the TED has a unique ability to spontaneously form an intramolecular thioester bond between the conserved cysteine and glutamine side chains (*15*, *17*). Given that a C426A mutation in Cpa-TED–which prevents thioester formation–results in a ∼75% reduction in adhesion to keratinocytes (*17*), it was postulated that TEDs covalently anchor bacteria to host cells by reacting their internal thioester with a nucleophile on the host surface (*15*, *17*). Drawing on an analogy to the mammalian complement proteins C3 and C4, which utilize similar Cys–Gln thioesters to form covalent linkages with their targets, investigators initially presumed TED-mediated binding to be irreversible (*15*).

A TED was subsequently identified within the fibronectin-binding protein SfbI (*15*), a GAS adhesin causing cell adhesion and invasion. Using mass spectrometry, this study demonstrated that the SfbI-TED and its target, fibrinogen, form a complex that is cross-linked via an intermolecular isopeptide bond. This crosslinking results from an acyl-transfer reaction, driven by the nucleophilic attack of an amine group from fibrinogen on the TED thioester. This mechanism was termed a “chemical harpoon” and was proposed that it facilitates irreversible bacterial attachment to the host. Notably, unlike complement proteins C3 and C4, which utilize thioester-mediated crosslinking to react with a broad range of nucleophiles, TEDs exhibit remarkable target specificity. Analysis of fibrinogen binding across ten distinct TEDs showed that only SfbI- and FbaB-TED, both encoded within the FCT region of the GAS genome (*18*), were capable of binding fibrinogen. Moreover, mass spectrometry revealed that SfbI-TED specifically reacts with Lys100 on the fibrinogen Aα chain, differentiating it from the other available lysine residues. This finding indicates that the “chemical harpoon” mechanism involves a targeted recognition mechanism rather than a random nucleophilic attack on a reactive thioester (*15*).

Recent studies challenged the prevailing view that TED-mediated chemistry is irreversible (*19*, *20*). Using single-molecule force spectroscopy, these studies demonstrated that the Cpa thioester can be cleaved by a small-molecule nucleophile (methylamine) and reforms upon removal of the nucleophile, provided that the mechanical load is sufficiently low to permit TED refolding. These findings suggest that thioester reactivity is gated by the structural integrity of the TED fold, leading to the proposal of a “smart covalent bond” model in which the lifetime of the covalent linkage is modulated by mechanical force. However, these experiments relied on small-molecule amines rather than on physiological protein targets. Consequently, it remains to be elucidated whether intermolecular crosslinking between TEDs and their native ligands is similarly reversible (*19*, *20*).

Using SfbI-TED as a model system, we analyzed the kinetic and thermodynamic mechanisms underlying target recognition and crosslinking. Biolayer interferometry revealed a multistep binding process: initial non-covalent recognition, followed by thioester-mediated formation of an intermolecular crosslink, which is in dynamic equilibrium with the reverse reaction. Furthermore, we discovered that SfbI-TED displayed pronounced pH-responsive behavior; specifically, the SfbI-TED/fibrinogen complex dissociated rapidly under mildly acidic conditions (∼pH 6.0). Similar pH-dependent properties were observed in other TED homologues, suggesting that pH responsiveness is a conserved feature of the TED family rather than a unique attribute of SfbI. These observations suggest that the ability to function as a pH-responsive adhesin module may have contributed to the widespread dissemination and evolutionary success of TEDs in Gram-positive bacteria.

## Results

### pH-dependent changes in thermal stability

To characterize SfbI-TED, we first examined the pH dependence of its thermal stability. Previous work on Cpa indicated that the intramolecular thioester bond contributes to adhesion, but not to thermal stability; the melting temperature (*T*_m_) did not change in a thioester-deficient mutant (*21*). Therefore, we compared wild-type SfbI-TED with thioester-deficient mutants (Q261A, C109S, and C109A). Differential scanning calorimetry (DSC) revealed a marked pH dependence exclusively for the wild type (Fig. 1, A and B). At pH 7.3, the *T*_m_ of the wild type (52.10 °C) was comparable to that of the Q261A mutant (53.44 °C) (Table 1), which lacks the thioester bond, consistent with the previous report. However, as the pH decreased, an additional denaturation peak appeared around 65 °C specifically in the wild type; this higher-temperature peak became dominant as the pH declined. We attribute this behavior to the coexistence of molecules with an intact and cleaved thioester bond within the wild-type sample; the former gives rise to a higher-temperature peak and the latter to a lower-temperature peak. This interpretation is reinforced by the observation that the *T*_m_ values of all thioester-deficient mutants correspond to the lower-temperature peak of the wild type at each pH.

**Fig. 1.**
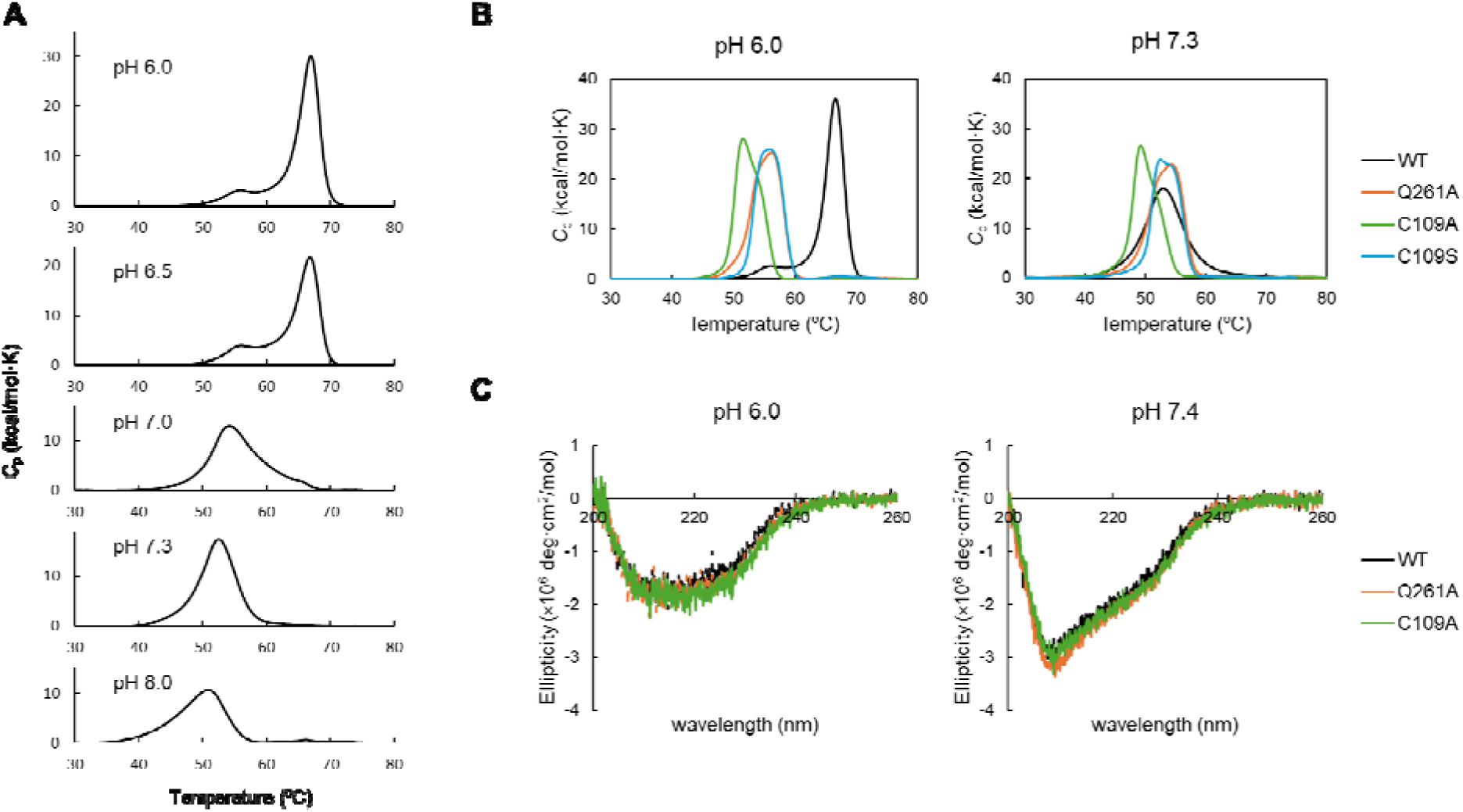
pH-dependent structural properties of SfbI-TED. (**A**) Thermal stability of SfbI-TED analyzed by DSC. The thermograms show the molar heat capacity (*C*_p_) as a function of temperature across a pH range of 6.0–8.0. (**B**) Comparison of DSC profiles between wild-type SfbI-TED and thioester-deficient mutants (C109A, C109S, and Q261A) at pH 6.0 and 7.3. (**C**) Far-UV CD spectra of SfbI-TED at pH 6.0 and 7.4.

**Table 1.** *T*_m_ of lower-temperature or higher-temperature peak.

| | | $T_m$ 1 (°C) | $T_m$ 2 (°C) |
| --- | --- | --- | --- |
| WT | pH 6.0 | 56.69 ± 0.39 | 66.54 ± 0.09 |
|  | pH 6.5 | 56.97 ± 0.12 | 66.44 ± 0.02 |
|  | pH 7.0 | 54.70 ± 0.13 | — |
|  | pH 7.3 | 52.10 ± 0.41 | — |
|  | pH 8.0 | 49.84 ± 0.35 | — |
| C109S | pH 6.0 | 55.76 ± 0.02 | — |
|  | pH 7.3 | 53.38 ± 0.04 | — |
| Q261A | pH 6.0 | 55.68 ± 0.23 | — |
|  | pH 7.3 | 53.44 ± 0.09 | — |
| C109A | pH 6.0 | 52.38 | — |
|  | pH 7.3 | 49.89 | — |

Data are shown as mean ± SD, n=3 (WT, C109S, Q261A), n=1 (C109A)

In all samples, including the mutants, thermograms shifted by about 2–5 °C between pH 6.0 and 7.3, indicating a small pH-dependent fluctuation even in the absence of the thioester bond (Table 1). To investigate whether pH affects structural conformation or packing, we measured the circular dichroism (CD) spectra at pH 6.0 and 7.4 (Fig. 1C). The spectra varied between pH conditions, primarily in the magnitude of the minimum at 208 nm. However, no clear change was observed in the ellipticity at 222 nm (reflecting α-helical content), or at 217 nm (reflecting β-sheet content) (*22*). The ellipticity at 208 nm is highly sensitive to the relative orientation of the helices (*23*, *24*), suggesting that SfbI-TED undergoes minor pH-dependent changes in the packing of its helical subdomain. This observation is consistent with the small (2–5 °C) shift in the DSC thermograms seen for all mutants, independent of the thioester bond (Fig. 1, A and B).

### pH-dependent cleavage and reformation of the thioester bond

To determine whether molecules with cleaved and intact thioester bonds coexist and whether their ratio changes with pH, we quantified free thiols using 4,4’-dithiodipyridine (4-PDS). This reagent reacts specifically with thiols to release 4-thiopyridone (4-TP; *ε*_324_ = 21,400 M^-1^ cm^-1^) (*25*) (Fig. 2A). Although the reaction between low-molecular-weight thiols and 4-PDS is typically rapid, we observed it to be extremely slow with folded SfbI-TED (fig. S1), suggesting that steric hindrance restricts the accessibility of 4-PDS. Consequently, we performed the reaction under denaturing conditions (Fig. 2B). We mixed SfbI-TED and sufficient 4-PDS at room temperature and rapidly heated to denature the TED fold. After rapid cooling and removal of aggregates, the absorbance at 324 nm (*A*_324_) was measured. Given that the side chain of C109 is the sole thiol in SfbI-TED, all thiols quantified in this assay were attributed to thioester bond cleavage. Normalized *A*_324_ (%) was calculated by dividing the observed *A*_324_ by the value expected for complete thioester-bond cleavage in 30 μM SfbI-TED.

**Fig. 2.**
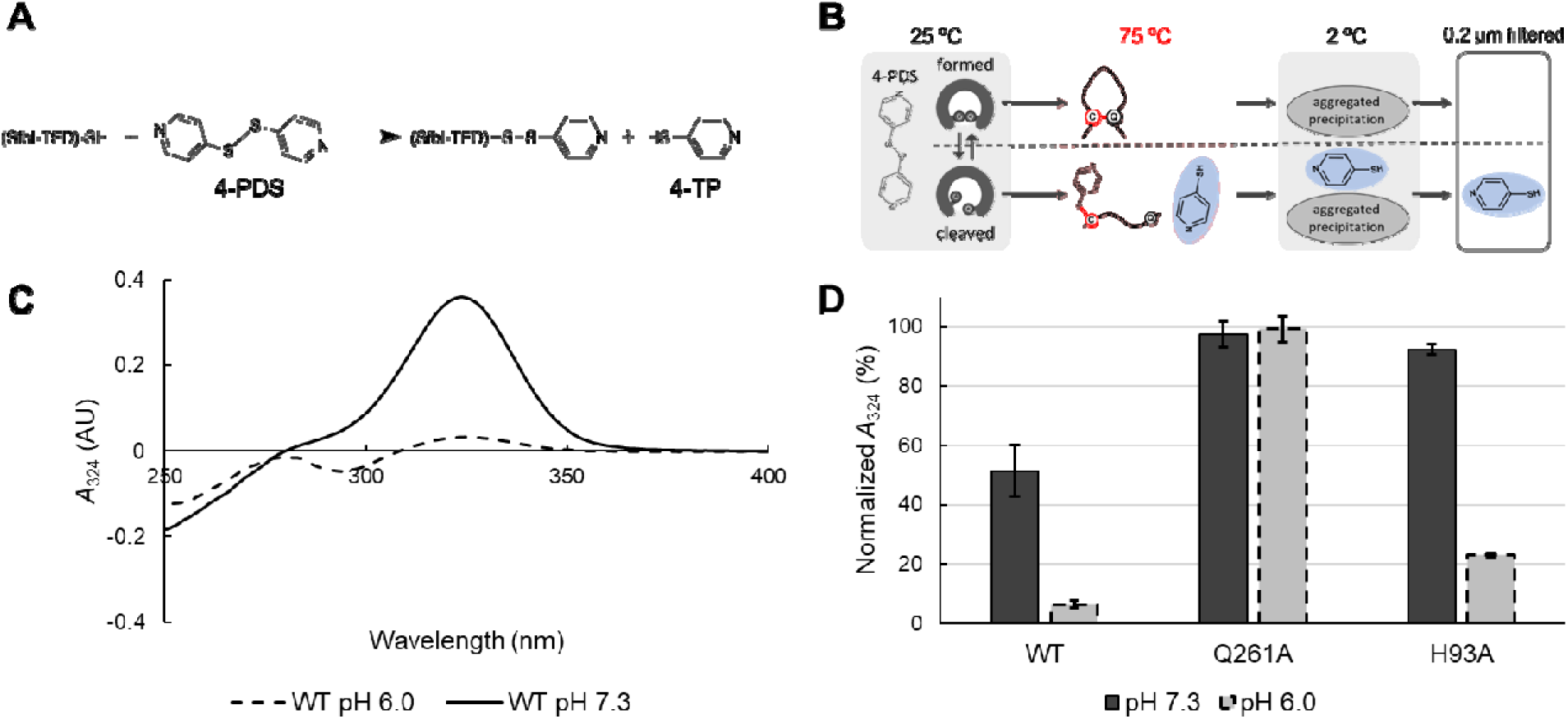
Quantification of intramolecular thioester bond cleavage. (**A**) Reaction scheme of thiol quantification using 4-PDS. 4-PDS reacts specifically with free thiols to release 4-TP, which absorbs light at 324 nm. (**B**) Schematic of the denaturation-based thiol quantification assay. SfbI-TED was incubated with 4-PD, heat-denatured to expose an internal thiol from a cleaved thioester, and rapidly cooled. Aggregates were removed by filtration prior to absorbance measurement. (**C**) Absorbance spectra of the reaction mixtures at pH 6.0 and 7.3 under denaturing conditions. (**D**) pH-dependent cleavage of the thioester bond. Normalized *A*_324_ represents the fraction of molecules with a cleaved thioester bond relative to the theoretical maximum (calculated for 30 µM SfbI-TED). Data are presented as mean ± s.d. (n=3).

Given that the thioester bond is generally stable under acidic and neutral conditions (with a hydrolysis half-life of ∼20,000 h at pH 7.0 (*26*)), the observed *A*_324_ values are most consistent with the thioester bond being pre-cleaved within the folded state.

As expected, the Q261A mutant, which lacks a thioester bond, showed nearly 100% normalized *A*_324_ at both pH 6.0 and 7.3. In contrast, the wild type exhibited marked pH dependence, with normalized *A*_324_ values being 6.5 ± 1.1% cmean ± SD, *n* = 3) at pH 6.0 and 51.5 ± 8.8% at pH 7.3 (Fig. 2, C and D). Importantly, this pH dependence persisted even after repeated dialysis between pH 6.0 and 7.3 (fig. S2), demonstrating a reversible equilibrium between the cleaved and intact states of the intramolecular thioester bond.

The finding that thioester bond cleavage is promoted at higher pH is consistent with the DSC results. The apparent discrepancy between the ∼50% cleavage at pH 7.3 observed in the thiol assay (Fig. 2D) and the single lower-temperature DSC peak at pH 7.3 (Fig. 1A) can be reconciled by the difference in heating rates; the thiol assay employs a rapid ramp (210 °C /min), whereas DSC uses a slow ramp (1 °C /min). In the 50–60 °C range, molecules with intact thioester bonds remained folded, whereas those with cleaved thioester bonds rapidly unfolded (Fig. 1A) and were therefore removed from the equilibrium between cleavage and reformation. Consequently, slower heating in the DSC allowed more extensive thioester bond cleavage to occur during the transition. The broader width of the lower-temperature DSC peak relative to the higher-temperature peak was also consistent with a Poisson-like process in which thioester bond cleavage is the rate-limiting event for unfolding.

In our experimental system, low-molecular-weight amines were absent, making hydrolysis a plausible mechanism for thioester cleavage. Indeed, mass spectrometry detected a peptide containing a Q261E conversion, which is indicative of thioester bond hydrolysis (fig. S3). Although only a single peptide containing Q261E was detected in our experiment, its strong negative charge (four acidic and one basic residue) likely reduced the ionization efficiency in the nano LC-MS/MS measurements. Collectively, these results support the occurrence of hydrolytic thioester bond cleavage.

The hypothesis that histidine protonation drives pH-dependent behavior was tested using the H93A mutant, in which the sole histidine in SfbI-TED is substituted with alanine. Thiol quantification showed a pH dependence similar to that of the wild type (Fig. 2D), indicating that H93 protonation was not involved.

### Model of SfbI-TED binding to fibrinogen

Using mass spectrometry, a previous study demonstrated that SfbI-TED binds specifically to Lys100 on the fibrinogen Aα chain (*15*). Given that the reaction between a thioester bond and a lysine side-chain amine is, in principle, a generic nucleophilic acyl transfer, such specificity is likely to stem from an additional mechanism: a protein–protein interaction that recognizes the local environment of Lys100. To account for this, we hypothesized a two-step reaction model wherein the surface residues of SfbI-TED first recognize the local environment of Lys100 on fibrinogen through non-covalent interactions, followed by thioester-mediated crosslinking (Fig. 3A).

**Fig. 3.**
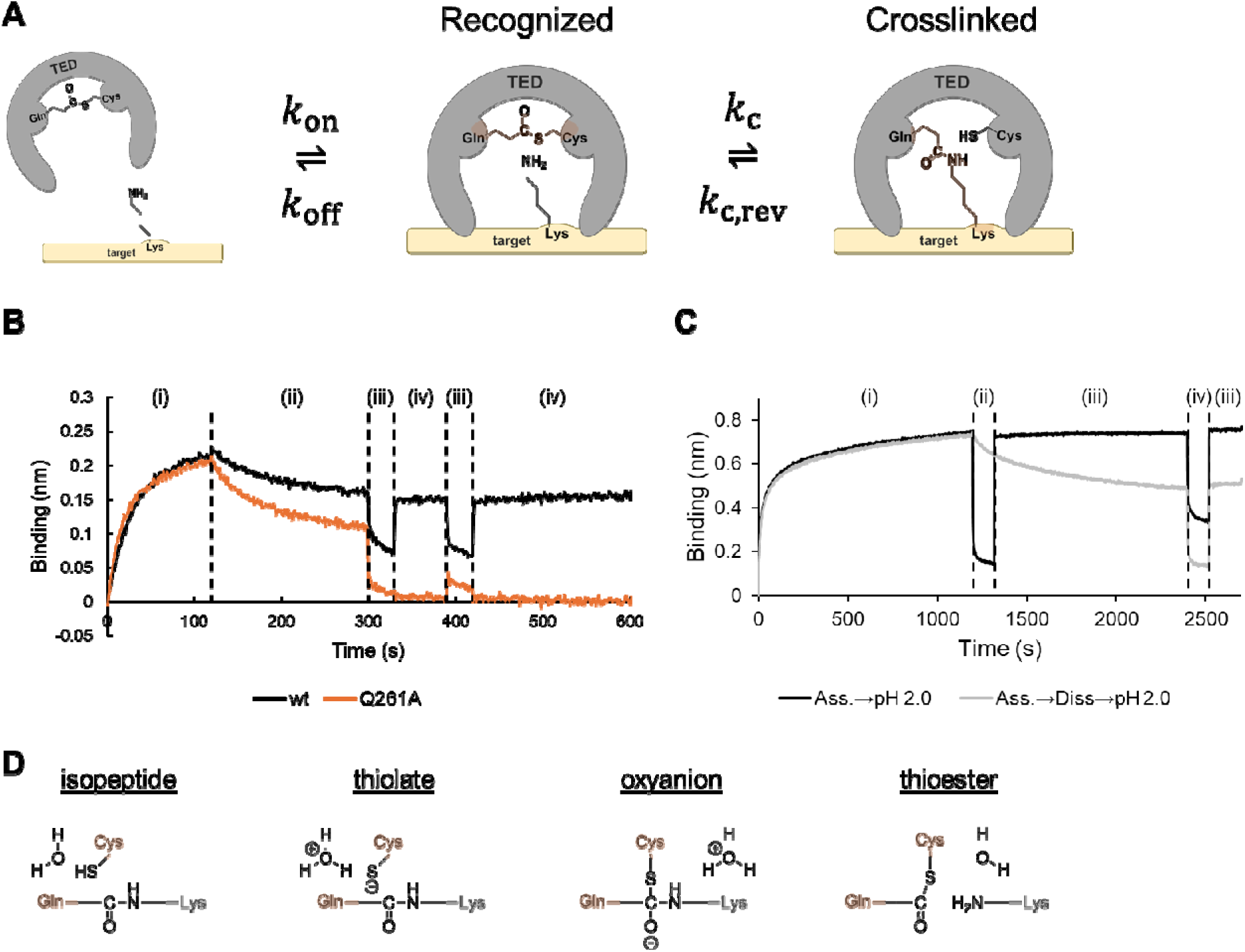
Reaction scheme of TED. (**A**) Reaction scheme of TED tested in this study, consisting of non-covalent target recognition (*k*_on_, *k*_off_), intermolecular crosslink formation (*k*_c_), and its reverse reaction (*k*_c,rev_). (**B**) BLI sensorgrams obtained with immobilized SfbI-TED. WT (black) or the crosslink-defective mutant Q261A (orange) was immobilized. (i) Association with 100 nM fibrinogen at pH 7.3; (ii) dissociation at pH 7.3; (iii) washing at pH 2.0; (iv) equilibration at pH 7.3. (**C**) BLI sensorgrams obtained with immobilized SfbI-TED WT. Two experimental sequences were compared; Association ➔ dissociation ➔ pH 2.0 wash (black) and association ➔ pH 2.0 treatment (gray). (i) Association with 5 *μ*M fibrinogen at pH 7.3; (ii) pH 2.0 (black) or pH 7.3 (gray); (iii) pH 7.3; (iv) pH 2.0. (**D)** Schematic of the proposed cleavage reaction of the intermolecular isopeptide bond formed between SfbI-TED (orange) and fibrinogen (gray).

Walden *et al.* suggested that the crosslinking may be irreversible, whereas Echelman and Alonso-Caballero et al. *et al.* showed—using methylamine as a surrogate target—that the crosslink can be cleaved (*15*, *19*, *20*). This reversibility is consistent with the spontaneous formation of the thioester bond within the TED, which is thought to proceed via nucleophilic attack of the thiol group of the Cys side chain on the carboxamide group of the Gln side chain. These findings suggest that cleavage of the intermolecular crosslink may also occur within the physiological complex. Based on these considerations, we tested the hypothetical reaction scheme shown in Fig. 3A.

To validate this model, we measured the interaction between fibrinogen and SfbI-TED using bio-layer interferometry (BLI). SfbI-TED was immobilized on the biosensor, which was subsequently immersed in fibrinogen solutions to monitor binding. The C109A mutant showed no detectable binding, whereas the Q261A mutant exhibited a distinct interaction with fibrinogen (fig. S4). This interaction demonstrates that SfbI-TED retains the capacity for non-covalent recognition of fibrinogen even in the absence of an intact thioester bond, validating that the model comprises at least two steps for the overall adhesion process (Fig. 3A).

Therefore, in this reaction model, the BLI response comprises contributions from both non-covalent and covalent species. To distinguish between these components, we applied a strong wash (pH 2.0) after the dissociation step (Fig. 3, B and C). Under this condition, the response of the Q261A mutant, which is unable to form an intermolecular crosslink, was completely abolished, whereas the signal for the wild type remained largely unchanged. These results indicate that the residual response after the pH 2.0 wash is attributable to the covalent crosslinked complex. For the wild type in Fig. 3B, a gradual decay in response was observed during the dissociation step. To determine whether this represents the dissociation of the non-covalent complex or the cleavage of the pre-formed intermolecular crosslinks, we compared two conditions: applying a pH 2.0 wash immediately after a 20-minute association with 5 µM fibrinogen, versus applying it after a 20-minute dissociation in running buffer (Fig. 3C). The reduction in the acid-resistant signal after the running-buffer dissociation step indicates that the covalent crosslink undergoes reversible cleavage under the dissociation condition (Fig. 3C), consistent with the reaction model shown in Fig. 3A.

The dissociation of this covalent complex is likely driven by the reversal of the crosslinking reaction; the intermolecular isopeptide bond undergoes nucleophilic attack by the cysteine side chain of TED, resulting in bond cleavage and reformation of the intramolecular thioester bond within the TED (Fig. 3D). Force spectroscopy studies demonstrated that the thioester bond of Cpa-TED loses its reactivity toward methylamine when TED is unfolded under mechanical force (*19*, *20*). These findings indicate that both the acyl-transfer reaction and its reversal (Fig. 3D) are catalyzed by structural features intrinsic to the TED fold.

### Dissociation of the crosslinked complex at mildly acidic pH

Based on our finding that the intramolecular thioester bond of SfbI-TED undergoes pH-dependent cleavage, we examined whether its interaction with fibrinogen was pH-dependent. Following the association step at pH 7.3, the sensor was transferred into dissociation buffers with varying pH values (Fig. 4, A and B). The Q261A mutant, which cannot form an intermolecular crosslink, showed no dependence on pH during dissociation, whereas the wild-type complex dissociated faster at lower pH. These results indicate that non-covalent interactions are insensitive to pH, whereas the equilibrium governing the formation and cleavage of the intermolecular crosslink is strongly influenced by pH. The increase in the dissociation rate under mildly acidic conditions plateaued around pH 5.5 (Fig. 4B). Interestingly, the pH range over which the dissociation rate changed markedly (pH 5.5–8.0) coincides with the pH range of the GAS infection environment (pH 5.8–7.4) (*1–3*).

**Fig. 4.**
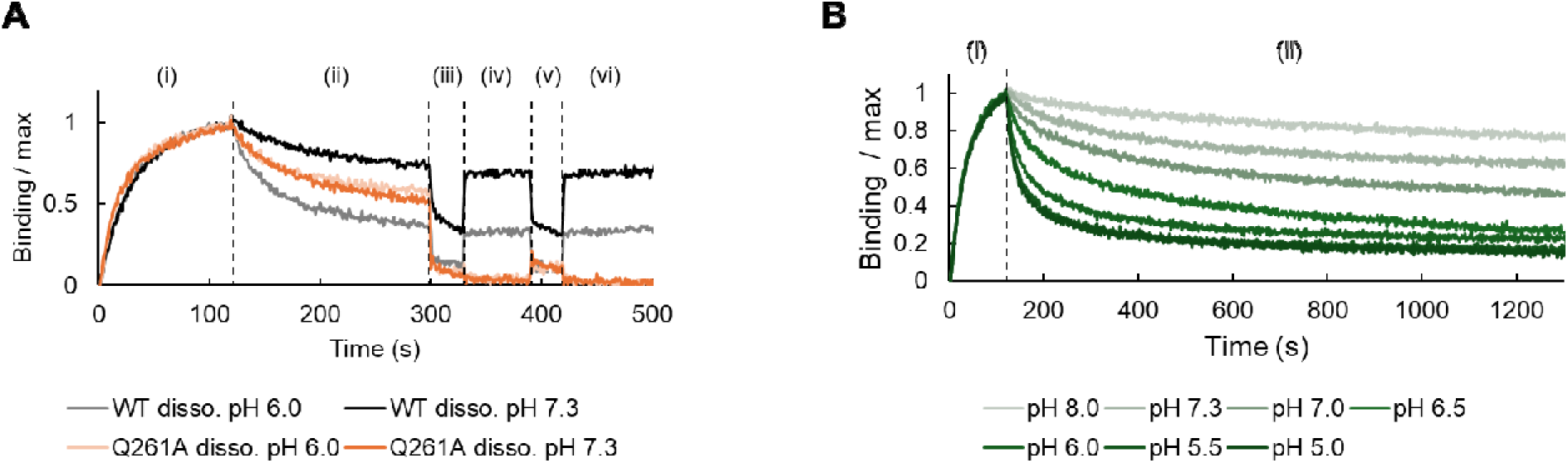
Kinetic mechanism of pH-dependent covalent binding. (**A**) Validation of covalent crosslinking by BLI. The sensorgrams display the sequential kinetic steps consisting of (i) association with fibrinogen in pH 7.3, (ii) dissociation in either pH 7.3 or pH 6.0, (iii)(v) strong wash in pH 2.0, and (iv)(vi) neutralization in pH 7.3. The vertical axis represents the response normalized to the maximum value in the association step. The pH 7.3 traces in Fig. 4A are replotted from Fig. 3B for direct comparison. (**B**) pH-dependent dissociation profiles. The sensorgrams display the kinetic response consisting of (i) an association phase with fibrinogen at pH 7.3, followed by (ii) a dissociation phase triggered by transferring the sensors into running buffers ranging from pH 5.0 to 8.0.

Furthermore, the observation that SfbI-TED, following the dissociation of fibrinogen at pH 6.0, could repeatedly form a covalently crosslinked complex upon rebinding to fibrinogen (fig. S4B) is consistent with our reaction model, wherein cleavage of the intermolecular crosslink is coupled with reformation of the intramolecular thioester bond (Fig. 3D).

### Identification of SfbI-TED residues involved in target recognition

Previous mass-spectrometric analysis established that SfbI-TED forms an intermolecular crosslink with Lys100 on the fibrinogen Aα chain(*15*). To gain structural insight into the binding pose, we predicted the complex structure using AlphaFold 3, incorporating the coiled-coil region of fibrinogen encompassing Lys100(*27*). Across 500 structures generated from 100 random seeds, the interface predicted Template Modeling (ipTM) scores were generally low (fig. S5); however, 260 of the 500 models placed SfbI-TED in close vicinity to Lys100. In these 260 models, the average distance between the carbonyl carbon atom of the TED thioester and the nitrogen atom of the Lys100 side-chain amine was 3.4 Å, with minimum distance of 2.3 Å (the model exhibiting the closest proximity between the reactive groups is shown in Fig. 5A).

**Fig. 5.**
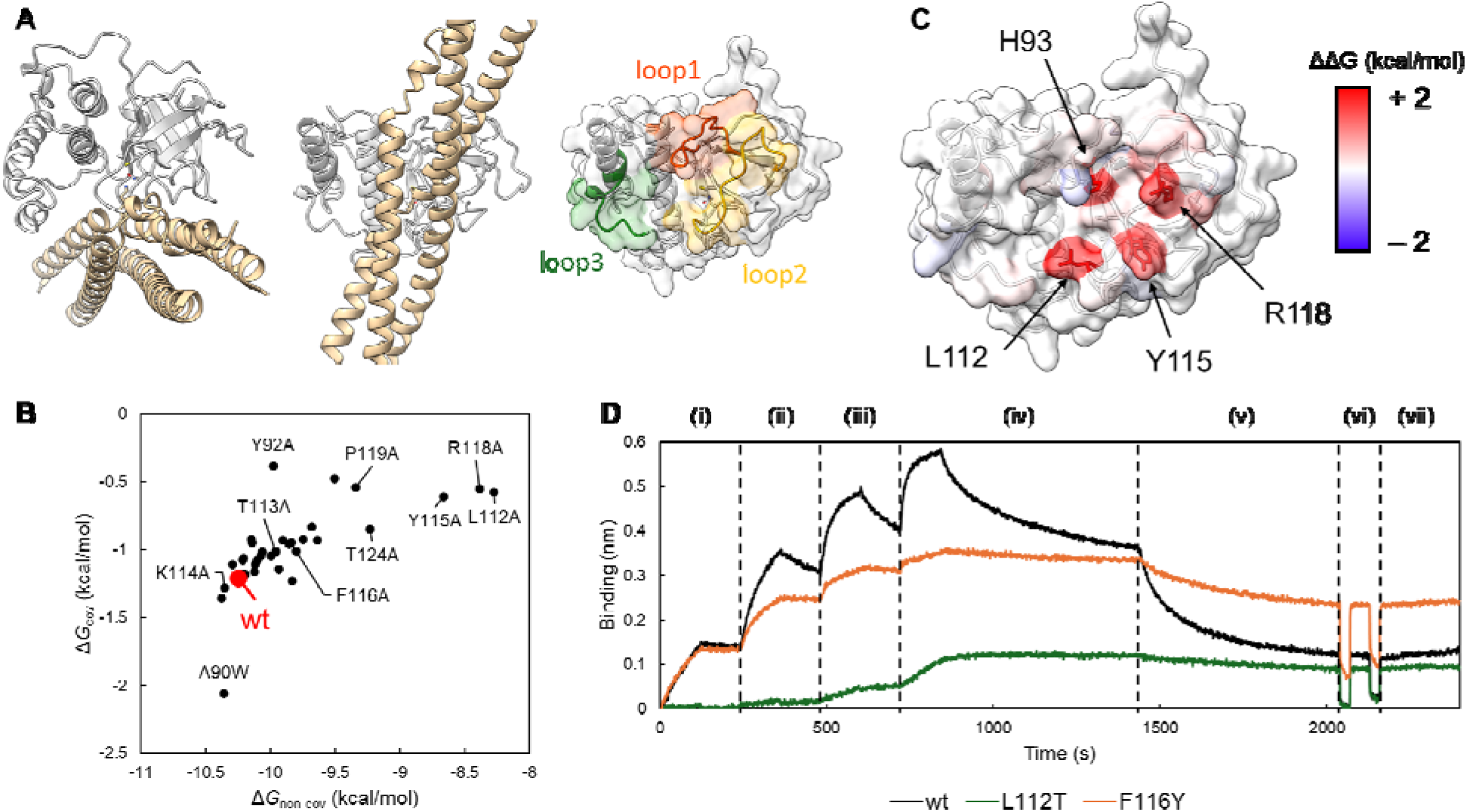
Residue-level contributions to SfbI-TED target recognition. (**A**) Structural model of the SfbI-TED–fibrinogen complex predicted by AlphaFold 3. SfbI-TED and fibrinogen are colored white and ochre, respectively. The side chains of C109 and Q261, which form the thioester bond, and the target lysine (K100) on fibrinogen are shown as sticks. The right panel highlights loop 1–3 at the putative binding interface, which were subjected to single-mutant analysis. (**B**) Correlation between the free energies of non-covalent association (-) and covalent crosslinking (). Data points represent individual mutants, revealing a positive correlation (Pearson’s *r* = 0.67). (**C**) Surface mapping of energetic contributions to non-covalent binding. Residues are colored according to the change in non-covalent binding free energy (-) upon alanine substitution, mapped onto the crystal structure of SfbI-TED (PDB 5A0L). Mutation of H93 (loop 1) abolished binding, identifying it as a key hotspot. L112, Y115, and R118 (loop 2) also contributed significantly, whereas loop 3 showed a negligible contribution. (**D**) Representative BLI sensorgrams for wild-type SfbI-TED and selected mutants (L112T and F116Y). The sensorgrams display the kinetic sequence consisting of (i)-(iv) association and dissociation steps at pH 7.3 across fibrinogen concentrations of 25, 100, 400, and 1600 nM, respectively, (v) a dissociation step at pH 6.0, (vi) two wash cycles in pH 2.0 buffer where all non-covalent interactions are dissociated, and (vii) the final response at pH 7.3, corresponding to the fraction of covalently crosslinked complex formed.

In the generated structures, the three loops of SfbI-TED formed a binding interface with fibrinogen. To experimentally validate their contribution, we performed a high-throughput single-mutant analysis using the FASTIA method(*28*). We targeted the 37 residues located on these loops. For the construct design, A90 was mutated to Trp, and owing to the failure to express D83A, the D83N mutant was substituted. Each mutant was immobilized on a BLI sensor and sequentially exposed to fibrinogen at 25, 100, 400, and 1600 nM (pH 7.3). The resulting sensorgrams were globally fitted to the two-step model to obtain kinetic parameters (*k_on_, k_off_, k_c_, k_c,rev_*) (fig. S6, S7). Notably, the H93A mutant did not bind. Although the H93F mutant was constructed subsequently, it failed to bind in a similar manner.

From the kinetic parameters obtained by fitting, the reaction free energies (Δ*G*) for each step of the binding model were calculated (Fig. 5B), and the change in the free energy of the non-covalent interaction (ΔΔ*G_non-covalent_*) was mapped onto the surface of SfbI-TED (Fig. 5C). Red residues corresponded to mutations that decrease the affinity, indicating a strong contribution of native side chains to target recognition. Only four residues showed a reduction in affinity corresponding to a ΔΔ*G_non-covalent_* of more than 1 kcal/mol, and complete loss of binding was observed exclusively for the H93A/F mutations (fig. S6). Thus, non-covalent target recognition by SfbI-TED was primarily mediated by the hotspot residue H93 (ΔΔ*G_non-covalent_* > 2 kcal/mol)(29), together with auxiliary interactions contributed by L112, Y115, and R118, which cluster around H93. In contrast, loop 3 was not involved in fibrinogen binding. Loops 1 and 2 flank the intramolecular thioester bond and their dominant contribution to fibrinogen recognition is consistent with the structural arrangement of the domain.

A notable intrinsic property of TED is the positive correlation between the free energies of the two-step reactions, with a Pearson correlation coefficient of 0.67 (Fig. 5B; see Methods and fig. S8 for the assessment). This result suggests that stabilization of the covalent complex necessitates the formation of a sufficiently stable non-covalent complex. This phenomenon is consistent with previous studies showing that TEDs can spontaneously reform their intramolecular thioester bonds, facilitating disengagement from unproductive reactions with small amines that are abundant *in vivo*(*19*, *20*, *30*).

During the course of our single-mutant analysis, we identified two mutants, F116Y and L112T, which bound more strongly than the wild type (Fig. 5D). Both mutants markedly stabilized the covalent complex, with L112T exhibiting almost irreversible binding at pH 7.3. Although these mutations also slowed dissociation at pH 6.0, they retained the characteristic pH dependence observed in the wild type, wherein dissociation was accelerated under acidic conditions compared to pH 7.3.

### Evidence for conserved pH-responsive behavior within the TED family

Finally, we assessed whether pH responsiveness is unique to SfbI-TED or is shared more broadly among TEDs. To address this, we examined the TEDs from two additional proteins: CpTIE-TED from *Clostridium perfringens* and FbaB-TED from GAS. For CpTIE-TED, the pH-dependent cleavage of the intramolecular thioester bond was measured using the same 4-PDS assay employed for SfbI-TED. Given that the only thiol-containing residue in CpTIE-TED is cysteine involved in the thioester bond, all detected thiols directly reflect bond cleavage. Consistent with SfbI-TED, CpTIE-TED displayed a clear pH dependence of thioester bond cleavage (Fig. 6A).

**Fig. 6.**
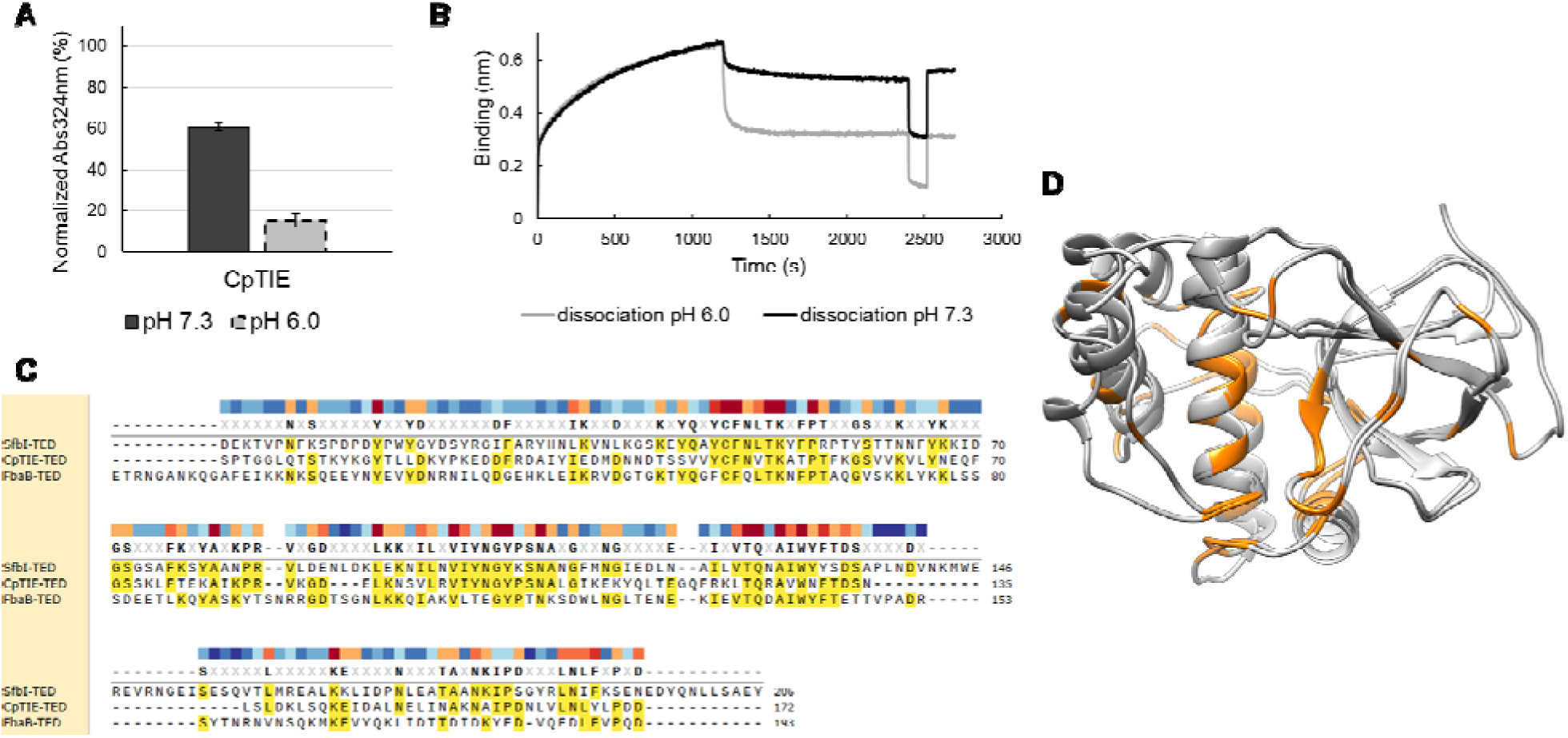
pH responsiveness in other TEDs. (**A**) pH-dependent reactivity of the intramolecular thioester bond in CpTIE-TED from *C. perfringens*. (**B**) BLI analysis of fibrinogen binding by FbaB-TED from GAS. The sensorgrams display kinetic phases consisting of association at pH 7.3, followed by dissociation in wash buffers at either pH 7.3 or pH 6.0, confirming that rapid acid-induced dissociation is a conserved feature. (**C**) Sequence conservation among TEDs. Pairwise sequence identities of CpTIE-TED and FbaB-TED relative to SfbI-TED are 26% and 25%, respectively. Residues conserved across two or three sequences are highlighted in yellow. The heat map above the alignment indicates conservation scores calculated by the Valdar method and visualized using SnapGene Viewer. (**D**) Structural alignment of SfbI-TED (PDB 5A0L) and CpTIE-TED (PDB 5A0G). Residues identical in sequence are colored orange.

FbaB is a surface protein encoded in the same FCT region of the GAS genome as SfbI(*18*). FbaB-TED also binds fibrinogen and possesses a C-terminal fibronectin-binding repeat region, rendering it similar to SfbI(*15*). Although FbaB is therefore not ideal for probing the evolutionary conservation of pH responsiveness across distantly related TEDs, it is, alongside SfbI, one of only two TEDs for which a physiological target has been identified. This renders it suitable for testing pH-dependent target binding. Using BLI, we found that FbaB-TED complexes with fibrinogen dissociated rapidly at pH 6.0 (Fig. 6B).

The sequence identity between CpTIE-TED and SfbI-TED was only 26%, while that between FbaB-TED and SfbI-TED was 25%, with most of the conserved positions confined to the buried core (Fig. 6, C and D). The observation of consistent pH responsiveness despite such low sequence identity suggests that this behavior arises from features intrinsic to the TED fold rather than from sequence-specific determinants.

## Discussion

In this study, we demonstrated that the interaction between SfbI-TED and fibrinogen is a reversible two-step process composed of non-covalent recognition followed by intermolecular isopeptide bond formation, with the latter being pH-dependent. In this section, we demonstrate that this pH responsiveness is rooted in the intrinsic pH dependence of thioester chemistry that is conserved across the TED family. Based on these findings, we propose a new hypothesis regarding the physiological function of TEDs, which serve as adhesins widely distributed among Gram-positive bacteria.

To formalize the pH dependence of the reaction free energy, we employed Alberty’s framework of transformed Gibbs energies, defined at a fixed pH, with all other species in their standard states(*31*). Within this framework, the reaction free energy can be expressed as the sum of a pH-independent term (Δ_r_*G*°) and a pH-dependent term (Fig. 7A). Accordingly, changes in the reaction free energy upon a pH shift are fully captured by the latter term, such that only ΔΔ_f_*G*′°cpH 6.0 -> 7.3) needs to be considered. ΔΔ_f_*G*′° can be further decomposed into two contributions (Fig. 7B). One term, 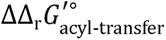, arises from protonation equilibria of the functional groups directly involved in the thioester reaction, namely the thioester bond itself and the nucleophile. The second term, 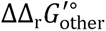encompasses all remaining contributions, including changes in the p*K*_a_ values of general acid–base catalytic residues surrounding the thioester bond. Crucially, the former contribution can be explicitly calculated within Alberty’s framework. Detailed theoretical discussions are provided in the Supplementary Text.

**Fig. 7.**
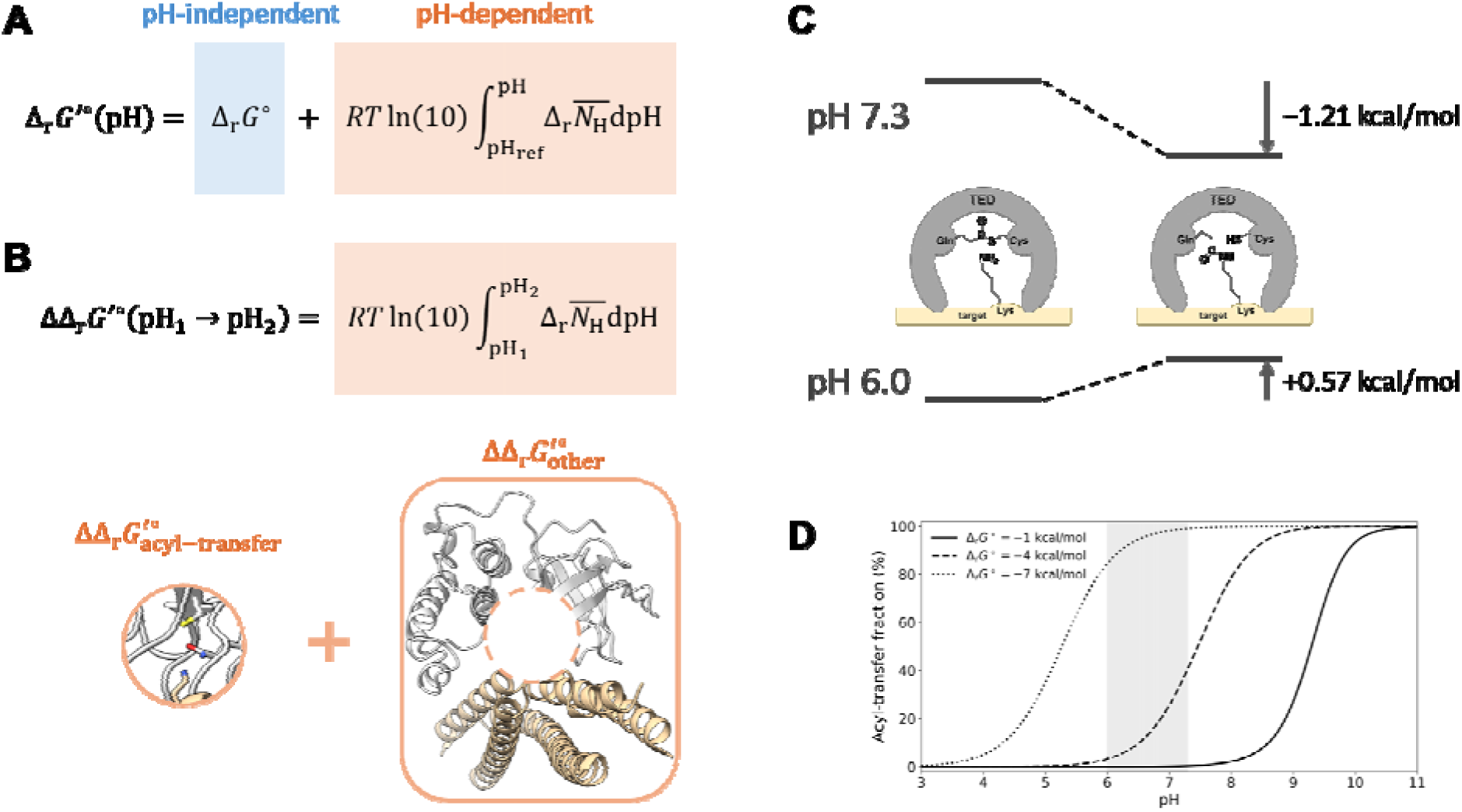
Graphical representation of the thermodynamic analysis of pH responsiveness. (**A**) Decomposition of transformed standard reaction free energy into pH-independent and pH-dependent terms according to Alberty’s framework. denotes the change in the average number of bound protons accompanying the reaction. (**B**) Decomposition of the pH-dependent change in the transformed standard reaction free energy into two contributions: one arising from the reactive triad, comprising the thioester-forming cysteine and glutamine as well as the target lysine, and the other encompassing all remaining factors. Structure adapted from Fig. 4a. (**C**) Thermodynamic switching of reaction favorability in the intermolecular isopeptide bond formation. (**D**) Effect of the pH-independent term on the manifestation of intrinsic pH responsiveness of the acyl-transfer reaction at the macromolecular level.

First, we analyzed the pH-dependent reaction free energy of the intramolecular thioester bond cleavage. A plausible mechanism for this cleavage reaction is hydrolysis, wherein new titratable groups emerge as the reaction proceeds:

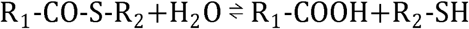

For this cleavage reaction, the theoretical value of 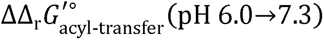 −1.77 (kcal/mol) (Eq. S6). This value can be perturbed as a function of the p*K*_a_ of the carboxyl group (R1-COOH) and the thiol group (R2-SH) and it is derived using typical solution values(*32*). Meanwhile, the pH responsiveness of the reaction can be experimentally determined to be ΔΔ_r_*G*′°cpH 6.0->7.3) = -1.63 ± 0.24 ckcal/mol) cmean ± SD n=3), using the normalized *A*_324_ values (Fig. 2D) to estimate the equilibrium constant (Eq. S7, S8). From the difference between the experimentally determined ΔΔ_r_*G*′°cpH 6.0->7.3) and the theoretically calculated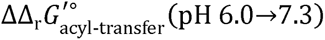, we obtain 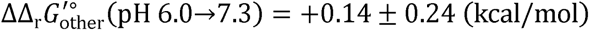 (Eq. S9). This result indicates that most of the macromolecular-level pH responsiveness of SfbI-TED originates from the acyl transfer reaction itself, with only minor contributions from the surrounding protein environment. Visual inspection of the region surrounding the thioester bond in the 3D structure of SfbI-TED did not reveal functional groups poised to act as general acid–base catalysts. Consistent with this observation, sequence conservation analysis across the Pfam-annotated TED family (PF08341) showed no conserved residues of this type near the thioester bond (fig. S12).

In the above analysis, we assumed that the cleavage detected by the thiol quantification assay primarily reflected the hydrolysis of the intramolecular thioester bond. Nevertheless, even in the absence of low-molecular-weight amines, aminolysis mediated by intrinsic amino groups of SfbI-TED cannot be entirely excluded. Importantly, even under these circumstances, our thermodynamic conclusions remain robust (Eqs. S10–15).

Having established the general principles of TED reactivity, we now turn to the reaction of primary physiological relevance: the formation and cleavage of the intermolecular isopeptide crosslink with the target protein. This reaction can be readily explained using the same thermodynamic framework.

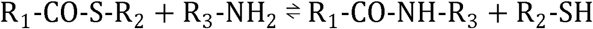

For this reaction as well, the theoretical value of 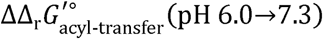 is -1.78 ckcal/mol), considering only the three residues that are directly involved in the acyl-transfer reaction; the pH responsiveness at the macromolecular level can be rationalized by additionally incorporating 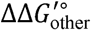 (Eq. S17–18). As in the case of intramolecular thioester bond cleavage, the contribution of 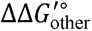 is unlikely to be sufficient to appreciably counterbalance the pH dependence dictated by 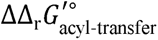 in the intermolecular reaction either. As an order-of-magnitude estimate, assuming that p*K*_a_ shifts within the complex arising from acyl-transfer are negligible, the reaction free energy for intermolecular crosslink formation is −1.21 kcal/mol (favorable) at pH 7.3, whereas it shifts to +0.57 kcal/mol (unfavorable) at pH 6.0, as calculated based on FASTIA results (Fig. 5B). This confirms a switch in thermodynamic favorability across this pH range (Fig. 7C).

Notably, these considerations are solely from an equilibrium perspective, which alone does not guarantee that the reaction will proceed at an appreciable rate. However, BLI analysis demonstrated that acidification accelerates dissociation (Fig. 4B). Overall, while the SfbI-TED–fibrinogen complex remains stable near neutral pH, it dissociates at a biologically relevant rate in response to environmental acidification.

Next, we investigated whether pH-responsive binding is conserved across the TED family. Since the conserved acyl transfer reaction itself intrinsically exhibits pH responsiveness, this regulation likely represents a general feature of TEDs. Notably, however, the reaction free energy consists of a pH-independent term (Δ_r_*G*°) and a pH-dependent term (Fig. 7A); whether this intrinsic pH responsiveness manifests at the macromolecular level depends on the balance between these two contributions.

For example, increasing the instability of the thioester bond promotes intermolecular crosslinking. Independently, strengthening the target-specific non-covalent interactions further facilitates this crosslinking (Fig. 5B). Under such conditions, the contribution of the pH-independent term (Δ_r_*G*°) dominates, requiring more acidic conditions for efficient cleavage reaction (Fig. 7D); consequently, the pH-responsive component may be effectively masked within the physiologically relevant pH range, precluding a detectable shift. Indeed, in the SfbI-TED L112T mutant, the fibrinogen complex remained stable across the tested pH range.

Notably, CpTIE-TED exhibits pH responsiveness despite its distant sequence and taxonomic relationship to SfbI-TED. Thiol quantification confirmed that the thermodynamic favorability of the CpTIE-TED reaction switches between pH 6.0 and 7.3 (Fig. 6A). Similarly, although the trait of near-irreversible binding to fibrinogen is accessible through a single L112T mutation, SfbI-TED likely avoided this course of evolution. These findings strengthen the hypothesis proposed by Echelman and Alonso-Caballero *et al.* that reversible adhesion confers an advantage to bacteria (*19*, *20*). Furthermore, excessive instability of the intramolecular thioester bond hinders efficient repair following cleavage by the abundant amines present *in vivo*(*30*). Taken together, these observations support the generalization that, in principle, pH-responsive adhesion represents an intrinsic property shared across the TED family, although this property may obscure this trait in some evolved TEDs.

Previous studies postulated that TEDs might function as molecular anchors that irreversibly tether bacteria to host tissues. In contrast, our results demonstrate that SfbI-TED binding is reversible, with a lifetime comparable to other adhesins. This raises a fundamental question: What biological advantage does covalent adhesion provide? While previous studies proposed that mechanical force modulates “smart covalent bond” lifetimes, we propose a distinct but complementary perspective: TEDs have proliferated because they act as versatile, mechanism-based, pH-responsive adhesion modules. In GAS, for example, fibrinogen binding is reported to sterically inhibit host phagocytosis(*33*), yet this same coating impairs the function of other adhesins(*34*). Consequently, SfbI’s ability to switch fibrinogen binding on and off in a pH-responsive manner confers a strategic advantage.

Conventional pH-responsive binders typically rely on interface residues whose protonation states vary with pH(*35*). In contrast, the thioester bond embedded within the structural core intrinsically confers pH responsiveness to TEDs. Consequently, TEDs retain pH-dependent binding regardless of the target-recognition interface diversity or target surface properties, provided the reaction utilizes thioester chemistry. We proposed that this modular ability to implement pH-responsive binding via a conserved catalytic motif, without constraining the evolving binding interface, underlies the widespread distribution and evolutionary success of TEDs across Gram-positive bacteria.

Research on bacterial adaptation to environmental pH has traditionally focused on pH-dependent gene expression. Although such transcriptional regulation drives infection dynamics and virulence, the pH modulation of protein function warrants equal consideration, as these mechanisms enable adaptive responses on different timescales. Our findings suggest that such protein-level regulation represents a widespread strategy for environmental adaptation in Gram-positive bacteria.

In this study, we combined *in vitro* characterization of physicochemical properties and interactions of TEDs with thermodynamic analysis to reveal that TEDs function as intrinsically pH-responsive adhesion modules, transcending species-specific regulation. These results suggest that pH responsiveness represents a unifying principle across the TED family. Despite these insights, several limitations remain. Our findings are primarily based on in vitro thermodynamic analyses, and how the complex microenvironment during infection—including molecular crowding and competing host factors—modulates this pH switch remains to be fully elucidated.

Future studies, including cellular and in vivo approaches, will be essential to verify how these modules detect environmental cues and influence physiological behavior during pathogenesis. Incorporating pH-dependent binding modulation into physiological analyses of TED family members will be critical for a comprehensive understanding of their biological roles.

## Materials and Methods

### Protein expression in *Escherichia coli*

With the exception of the mutants analyzed by FASTIA, all TEDs and SpyCatcher003(*36*, *37*) used in this study were expressed in *E. coli* BL21(DE3). Expression vectors were derived from pCold II (Takara Bio) for cytoplasmic expression. Wild-type and mutant SfbI-TED constructs contained N-terminal 6xHis-SpyTag003 tags. FbaB-TED was similarly tagged at the C-terminus. CpTIE-TED was tagged N-terminally with a 6xHis-SUMO. SpyCatcher003 was produced with either an N-terminal 6xHis alone or combined with C-terminal AviTag.

Plasmids were transformed into BL21(DE3) cells, and plated on LB agar containing 100 µg/mL ampicillin. Colonies were inoculated into 6 mL of LB broth containing 100 µg/mL ampicillin and cultured overnight at 37 °C with shaking at 200 rpm. The seed culture was then transferred to 800 mL of LB broth containing 100 µg/mL ampicillin in baffled flasks, which were incubated at 37 °C with shaking at 120 rpm. When the optical density at 600 nm (OD_600_) reached 0.4, the culture was cooled at 4 °C for 30 min. Protein expression was induced with 0.5 mM isopropyl β-D-1-thiogalactopyranoside (IPTG), followed by overnight incubation at 15 °C. Finally, cells were harvested by centrifugation (8,000 × g for 10 min) and the pellets stored till further processing.

### Purification by immobilized metal affinity chromatography (IMAC)

The cell pellet was resuspended in 30 mL of lysis buffer (20 mM Tris-HCl, 500 mM NaCl, and 5 mM imidazole, pH 8.0) and lysed by sonication on ice for 15 min. The lysate was clarified by centrifugation (40,000 × g for 1 h), and the supernatant was filtered (0.2-µm filter) before loading onto a gravity-flow column (15 mm diameter, Bio-Rad) packed with Ni-NTA agarose resin (QIAGEN). All subsequent steps were performed at 4 °C. The resin was washed with 15 column volumes (CV) of wash buffer (20 mM Tris-HCl, 500 mM NaCl, and 20 mM imidazole, pH 8.0). Finally, proteins were eluted with 5 CV elution buffer (20 mM Tris-HCl, 500 mM NaCl, and 200 mM imidazole, pH 8.0).

To remove the N-terminal 6xHis-SUMO tag from CpTIE-TED, the IMAC eluate was incubated overnight at 4 °C with in-house-produced Ulp1 protease while dialyzing against 50 volumes of cleavage buffer (20 mM Tris-HCl, 150 mM NaCl, pH 8.0). Following digestion, the imidazole concentration was adjusted to 5 mM, and the sample was incubated with the Ni-NTA resin for 1 h with gentle rotation. The mixture was then applied to an empty gravity-flow column, and the flow-through containing untagged CpTIE-TED was collected.

### Size-exclusion chromatography (SEC)

The IMAC eluate (or the flow-through fraction for CpTIE-TED) was dialyzed overnight against 50 volumes of SEC buffer (20 mM HEPES-NaOH, 150 mM NaCl, pH 7.3). The protein was further purified via SEC on a HiLoad 26/600 Superdex 75 pg column (Cytiva) connected to an ÄKTA prime system equilibrated with SEC buffer. Fractions containing monomeric protein were pooled for subsequent experiments.

### Buffer exchange for biochemical and biophysical analyses

SEC-purified proteins obtained at pH 7.3 were used directly for experiments performed at that pH. For other pH levels, proteins were dialyzed into the following target buffer systems: 20 mM HEPES-NaOH and 150 mM NaCl for pH 7.0 and 8.0, or 20 mM Bis-Tris-HCl and 150 mM NaCl for pH 6.0 and 6.5. For CD spectroscopy, proteins were exchanged into PBS (pH 7.4) to minimize background absorbance in the far-UV region (< 200 nm). Standard dialysis involved 50 volumes of the target buffer overnight at 4 °C. For DSC measurements, a rigorous two-step dialysis was conducted: initial dialysis for 4 h against 50 volumes of buffer, followed by overnight dialysis against fresh buffer. Protein concentrations were determined by measuring the absorbance at 280 nm (*A*_280_) using a NanoDrop One spectrophotometer (Thermo Fisher Scientific) and adjusted for each experiment.

### Differential scanning calorimetry (DSC)

Wild-type SfbI-TED was analyzed across five pH conditions (6.0, 6.5, 7.0, 7.3, and 8.0), whereas the C109A, C109S, and Q261A mutants were analyzed at pH 6.0 and 7.3. All samples were prepared at a concentration of 20 µM.

DSC measurements were performed using a MicroCal PEAQ-DSC automated system (Malvern Panalytical). Thermal scans were conducted from 20 °C to 110 °C at a scan rate of 1 °C/min. The data within the 30–80 °C range, which encompassed the observed thermal transitions, were utilized for analysis. Buffer baselines were subtracted, and the resulting thermograms were fitted to the Non-Two-State model using the MicroCal PEAQ-DSC Software (version 1.40).

### Circular dichroism (CD) spectroscopy

Far-UV CD spectra of wild-type SfbI-TED and the C109A and Q261A mutants were recorded at 25 °C (pH 6.0 and 7.4) using a J-1500 spectropolarimeter (JASCO). Samples (5 µM) were measured in a 1-mm path length quartz cuvette. Spectra were obtained by scanning from 260 to 200 nm at 60 nm/min, with five accumulations averaged per sample. Following buffer-baseline subtraction from the raw data, the corrected spectra were converted to molar ellipticity based on the protein concentration and path length.

### Thiol quantification upon thermal denaturation

Free thiol content was quantified using 4,4′-dithiodipyridine (4-PDS). A 100 mM stock solution of 4-PDS was prepared in methanol. Protein samples (SfbI-TED or CpTIE-TED) were prepared in either pH 7.3 buffer (20 mM HEPES-NaOH, 150 mM NaCl) or pH 6.0 buffer (20 mM Bis-Tris-HCl, 150 mM NaCl). The protein solutions and 4-PDS stock were equilibrated to room temperature and mixed to yield final concentrations of 30 µM protein and 100 µM 4-PDS, respectively, in a 200 µL volume. Control samples contained 100 µM 4-PDS in each corresponding buffer without the addition of protein.

Immediately after mixing, samples were transferred to a thermal cycler (MiniAmp Plus, Thermo Fisher Scientific), and subjected to the following thermal profile: heating from 25 °C to 75 °C at a ramp rate of 3.5 °C/s, holding at 75 °C for 30 s, cooling to 2 °C at a rate of 3.5 °C/s, and holding at 2 °C for 2 min. Following this treatment, all protein samples (wild-type SfbI-TED, mutants, and CpTIE-TED) exhibited visible turbidity. Aggregates were removed via filtration through 0.2-µm centrifugal devices (Nanosep, Pall Corporation) at 5,000 rpm for 1 min. The resulting filtrates were immediately analyzed by UV–visible spectroscopy.

Absorbance spectra were recorded from 400 to 240 nm using a V-660 spectrophotometer (JASCO) equipped with a water-cooled Peltier cell changer (PAC-743). An empty cuvette served as the baseline. To account for background absorbance and potential thermal degradation of 4-PDS, the spectrum of the protein-free control was subtracted from the corresponding protein-containing sample. Normalized absorbance at 324 nm (*A*_324_) was calculated as follows:

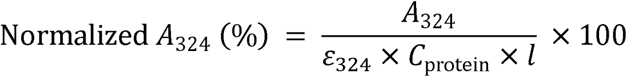

where ε_324_ is the extinction coefficient of 4-thiopyridone (4-TP) at 324 nm (21,400 M□¹ cm□¹, determined experimentally from its reaction with cysteine in a previous study(*25*)), *C*_protein_ is the molar concentration of the protein (30 µM), and *l* is the path length (1 cm).

### Reactivity of SfbI-TED with 4-PDS under native conditions

To monitor thioester-bond cleavage in folded SfbI-TED, 30 µM SfbI-TED (wild type or Q261A) was incubated with 100 µM 4-PDS in pH 7.3 buffer (20 mM HEPES-NaOH, 150 mM NaCl) or pH 6.0 buffer (20 mM Bis-Tris-HCl, 150 mM NaCl). *A*_324_ was recorded every 10 s for up to 24 h at room temperature. To exclude a direct reaction between 4-PDS and the thioester moiety, control reactions utilized 30 µM acetylthiocholine iodide as a small-molecule thioester model. Background absorbance changes were assessed by monitoring 100 µM 4-PDS in the respective buffers without protein or thioester compounds.

### Identification of peptides after thioester hydrolysis by nanoLC-MS/MS

SfbI-TED samples (pH 7.3) were mixed (1:3 volume ratio) with an SDS-PAGE loading buffer lacking amines and reducing agents (50 mM HEPES-NaOH, 18.4 g/L SDS, 4.5% glycerol, 0.013 g/L bromophenol blue, pH 7.3) and incubated at room temperature for 10 min. Samples were separated by SDS-PAGE in running buffer (3 g/L Tris, 14.4 g/L glycine, and 1 g/L SDS) at 200 V for 60 min. Following Coomassie Brilliant Blue staining, protein bands were excised, destained, and subjected to in-gel digestion with trypsin at 37 °C for 16 h. Proteome analysis was performed by Orbitrap Eclipse Tribrid mass spectrometer with a FAIMS Pro interface (Thermo Fisher Scientific, Waltham, MA) coupled to Vanquish Neo UHPLC system (Thermo Fisher Scientific, Waltham, MA). The peptides were separated on a reversed-phase column using a linear gradient of 2–24% acetonitrile in 0.1% formic acid at a flow rate of 300 nL/min. Full MS scans were acquired in the Orbitrap at a resolution of 120,000, followed by MS/MS scans in the ion trap using higher-energy collisional dissociation (HCD) with a normalized collision energy of 35% and a maximum injection time of 10 ms. Peptide identification was carried out by searching against the customized database derived from UniProt human reference proteome (UP000005640) supplemented with the amino acid sequence of SfbI-TED using the Sequest HT algorithm in Proteome Discoverer Software (version 2.5) (Thermo Fisher Scientific, Waltham, MA, USA).

### Biotinylation of SpyCatcher003

Purified SpyCatcher003 fused with an N-terminal 6xHis and a C-terminal AviTag (SpyCatcher003-His-Avi) was dialyzed overnight against 20 mM Tris-HCl and 50 mM NaCl at pH 8.0. Biotinylation was performed in a 500-µL reaction mixture containing 28 µM SpyCatcher003-His-Avi, 0.1 mg/mL BirA ligase, 50 mM Bicine, 10 mM MgSO , 10 mM ATP, and 50 µM D-biotin. The mixture was then incubated at 30 °C for 60 min. The reaction mixture was subsequently buffer-exchanged into 20 mM HEPES-NaOH and 150 mM NaCl (pH 7.3) using a PD-MiniTrap G-25 column (Cytiva).

### Bio-layer interferometry (BLI)

Bio-layer interferometry measurements were performed using an Octet RED384 system (Sartorius). All assays were conducted at 30 °C with plate agitation at 1,000 rpm. Unless otherwise specified, the running buffer contained 20 mM HEPES-NaOH, 150 mM NaCl, and 0.05% (v/v) Tween-20.

### Assay 1: Binding analysis (Fig. 3B and Fig. 4)

SAX2 biosensors (Sartorius) were hydrated for 10 min and loaded with 2 µM biotinylated SpyCatcher003-His-Avi for 5 min. To remove loosely bound proteins, the biosensors were regenerated using 10 mM glycine (pH 2.0). SfbI-TED (10 µM; wild type, C109A, or Q261A) was then captured for 20 min via the SpyTag003–SpyCatcher003 reaction(*38*). Following capture, a second wash with 10 mM glycine (pH 2.0) was performed. The establishment of a stable baseline after this wash confirmed the removal of unreacted SfbI-TED and formation of a covalent complex.

Analyte solutions were prepared by dissolving human fibrinogen (Sigma-Aldrich) in running buffer and filtering through a 0.2-µm Nanosep centrifugal device (Pall Corporation). The fibrinogen concentration was determined via *A*_280_. Association kinetics were measured by dipping the TED-immobilized biosensors into 100 nM fibrinogen, followed by dissociation in running buffer at the indicated pH values (pH 7.3 for Fig. 3B; pH 6.0 and 7.3 for Fig. 4A; pH 5.0, 5.5, 6.0, 6.5, 7.3, and 8.0 for Fig. 4B). After each dissociation, biosensors were regenerated with 10 mM glycine (pH 2.0) and re-equilibrated in running buffer. After re-equilibration, the association–dissociation cycle was repeated to confirm that fibrinogen-binding capacity was retained. The C109A mutant exhibited no detectable binding to fibrinogen and showed lower nonspecific adsorption than SpyCatcher003-only biosensors; therefore, the C109A signal was subtracted as a reference from the wild type and Q261A data.

### Assay 2: Covalent crosslinking analysis (Fig. 3C and Fig. 6B)

SpyCatcher003 fused with an N-terminal 6xHis (10 µM) was diluted 20-fold into 10 mM sodium acetate (pH 4.5) and immobilized onto AR2G biosensors (Sartorius) pre-activated with EDC/sulfo-NHS for 2 min. Unreacted esters were quenched with 1 M ethanolamine (pH 8.3). SfbI-TED or FbaB-TED was captured via the SpyTag003–SpyCatcher003 reaction for 30 min, followed by regeneration with 10 mM glycine (pH 2.0).

Fibrinogen was prepared as described above and used at a concentration of 5 µM. For SfbI-TED, two conditions were tested following fibrinogen binding at pH 7.3: immediate regeneration with 10 mM glycine (pH 2.0), or regeneration after a dissociation step at pH 7.3. For FbaB-TED, biosensors were allowed to dissociate at pH 7.3 or 6.0 prior to regeneration with 10 mM glycine (pH 2.0). As in Assay 1, the C109A mutant was used as a reference, and its signal was subtracted from those of the other sensorgrams.

### Complex structure prediction by AlphaFold 3

Fibrinogen is a hexamer composed of two sets of Aα, Bβ, and γ chains (PDB 3GHG)(*39*). For complex modeling, the sequences corresponding to the coiled-coil region of the fibrinogen heterotrimer were used, focusing on the region containing Lys100 of the Aα chain, which was previously identified as the SfbI-TED binding site(*15*).

Structure prediction was performed using the AlphaFold 3. Calculations were performed using 100 random seeds to sample the conformational space extensively and generate 500 predicted complex structures. For each model, the distance was calculated between the carbonyl carbon of the SfbI-TED thioester and the ε-amino nitrogen (NZ) of lysine residues on fibrinogen. Lysine residues with an interatomic distance of less than 5 Å were defined as being positioned for recognition by TED. The model exhibiting the shortest nucleophile–electrophile distance was selected for visualization.

### High-throughput mutational analysis using FASTIA

High-throughput mutational scanning was performed using the FASTIA platform(*28*). A pET28b-based plasmid encoding SfbI-TED fused to the N-terminal 6xHis-SpyTag003 served as a template for generating individual SfbI-TED mutants. Single mutations were introduced into the residues within the putative fibrinogen-binding interface, as predicted by AlphaFold 3.

Linear DNA fragments encoding each mutant were generated by inverse polymerase chain reaction (PCR) using KOD One DNA polymerase (TOYOBO) and mutagenic primers. The template plasmid was removed by DpnI digestion and the PCR products were purified using a NucleoSpin 8 PCR Clean-up kit (MACHEREY-NAGEL). The fragments were circularized using the NEBuilder HiFi DNA Assembly Kit (New England Biolabs). The resulting circular DNA was used as a template for a second PCR amplification with the primers PUREfrex-F and PUREfrex-R to generate linear expression templates containing the T7 promoter, ribosome-binding site, 6xHis, SpyTag003, and the SfbI-TED coding region. These linear DNA fragments were purified and sequenced by Sanger sequencing (Fasmac).

Cell-free protein synthesis was carried out using the PUREfrex 2.1 system (GeneFrontier). Reaction mixtures (10 µL) were supplemented with 4 mM reduced glutathione, 3 mM oxidized glutathione, 5 µM DnaK, 1 µM DnaJ, 1 µM GrpE, 1 µM DsbC, and 5 ng of template DNA. Reaction mixtures were incubated at 37 °C for 16 h. Following synthesis, the mixtures were diluted eight-fold in running buffer (20 mM HEPES-NaOH, 150 mM NaCl, 0.05% (v/v) Tween-20, pH 7.3) supplemented with 20 mM MgCl_2_ to stabilize ribosomes and used directly for BLI immobilization.

Interaction kinetics were analyzed using BLI on the SAX2 biosensors. SpyCatcher003-His-Avi was used without *in vitro* biotinylation, relying on partial biotinylation by endogenous ligases during expression in *E. coli*(*40*). This allowed immobilization at higher SpyCatcher003 concentrations than in the purified assay, yielding loading responses of approximately 0.5–0.6 nm. The immobilization protocol was identical to that described above, except that an acidic wash (10 mM glycine-HCl, pH 2.0) was omitted after capturing SfbI-TED through the SpyCatcher003-SpyTag003 reaction to prevent the denaturation of potentially unstable mutants. Stable baselines confirmed that most of the SfbI-TED mutants were successfully immobilized.

Kinetic parameters were determined using single-cycle kinetics. The biosensors were sequentially dipped into four increasing concentrations of fibrinogen (25, 100, 400, and 1600 nM) for 120 s in running buffer. Each association step was followed by a dissociation step in the running buffer for 120 s. After the final association step at 1600 nM, the dissociation was monitored for 10 min. Finally, the biosensors were transferred to dissociation buffer adjusted to pH 6.0, to assess the dissociation kinetics under acidic conditions. The H93F mutant exhibited no detectable binding to fibrinogen and was therefore used as a reference; its signal was subtracted from those of the other mutants. Measurements were performed in triplicate.

### Curve fitting for FASTIA mutant analysis

Equilibrium constants for the non-covalent interaction and intermolecular crosslinking were determined by fitting the series of association and dissociation curves at pH 7.3 to the two-step reaction model described in the main text. The parameter vector ***p*** was defined as follows:

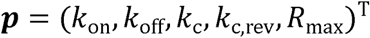

For each time point *t* = *t*_i_, the residual *r*_i_ was calculated as the difference between the measured response *R*(*t*_i_) and model response *R*_model_(*t*_i_; ***p***):

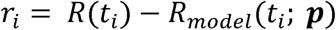

The residual vector ***r*** was defined as:

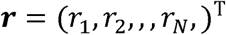

where *N* denotes the total number of data points in the time series recorded at 1 s intervals.

Parameter optimization was performed using the Trust Region Reflective algorithm implemented in the scipy.optimize.least_squares function (SciPy library) (*41*) to minimize the squared Euclidean norm of ***r***. Based on the optimized parameters, the equilibrium constants were calculated as:

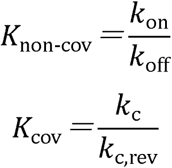

The corresponding reaction free energies, Δ*G_non-cov_* and Δ*G_cov_*, were calculated using the standard thermodynamic relationship:

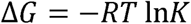

### Evaluation of parameter reliability

Since all fitting parameters—including those subsequently used to calculate Δ*G_non-cov_* and Δ*G_cov_* —were estimated simultaneously from a single sensorgram, a rigorous evaluation of parameter standard errors and identifiability was essential.

To evaluate the reliability of the derived free energies, we performed error-propagation analysis. After the optimization of the parameter vector ***p***, the Jacobian matrix ***J*** (N × P) was calculated as:

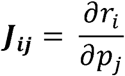

The parameter covariance matrix ***C_p_*** was estimated as:

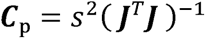

where *s*^2^ is the noise variance estimate calculated from the residual sum of squares:

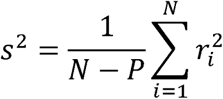

Here, *N* = 1,440 represents the total number of data points in the 1,440-second sensorgram (recorded at 1-s intervals), and *P* = 5 is the number of optimized parameters.

Next, the covariance matrix of the equilibrium constants (*C_K_*) was estimated by linear error propagation using the Jacobian of the equilibrium constants with respect to ***p*** (I_K_) as follows:

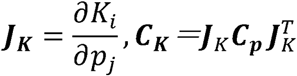

Similarly, the covariance matrix of the reaction free energies (*C*_Δ*G*_) was derived using the Jacobian of the free energies with respect to the equilibrium constants (***J***_Δ***G***_):

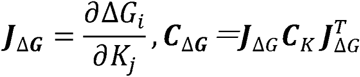

The standard errors of each *C*_Δ*G*_ were taken as the square roots of the diagonal elements of *C*_Δ*G*_. Fits were accepted only if they satisfied the criterion:

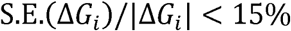

To further exclude fits where Δ*G_non-cov_* and Δ*G_cov_* were not independently identifiable, we applied the Motulsky and Christopoulos criterion(*42*), requiring that the magnitude of the error-correlation coefficient p_ΔG_ must not exceed 0.9:

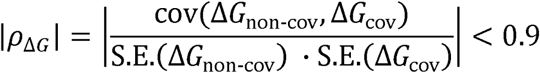

where cov(Δ*G_non-cov_*,Δ*G_cov_*) is the off-diagonal element of the 2 × 2 matrix *C*_Δ*G*_.

Only the fits satisfying both the relative standard error and correlation criteria were used for the final analysis. For each mutant, the mean values of Δ*G_non-cov_* and Δ*G_cov_* were calculated from the accepted fits. The Pearson correlation coefficient across mutants was determined to investigate the relationship between non-covalent interactions and intermolecular crosslinking reactions.

## Acknowledgments

We are grateful to GeneFrontier Corporation for providing the PUREfrex components.

## Funding

Japan Agency for Medical Research and Development (AMED), JP223fa627001 (UTOPIA), K T

Japan Agency for Medical Research and Development (AMED), JP223fa727002 (SCARDA), KT

Japan Agency for Medical Research and Development (AMED), JP24ama121033 (BINDS), KT JSPS KAKENHI, 22K21343, KT

JST ACT-X, JPMJAX222I, RM

## Author contributions

Conceptualization: YT, RM, KT

Formal Analysis: YT

Funding Acquisition: RM, KT

Investigation: YT

Methodology: RM, YT

Resources: HKH, MO, KT

Supervision: RM, KT

Visualization: YT

Writing – original draft: YT

Writing – review & editing: RM, HKH, MO, KT

## Data and materials availability

All data needed to evaluate the conclusions in the paper are present in the paper and/or the Supplementary Materials. Additional data related to this paper may be requested from the authors.

