## Supplementary Materials for "Chemically encoded pH-tunable covalent adhesion by a bacterial thioester domain"

**The PDF file includes:**

Supplementary Text
Figs. S1 to S12
Tables S1 to S3

### Supplementary Text

#### Free-energy analysis using Alberty's transformed standard energy

Under the constant temperature, pressure, and the amount of substance(*31*),

$$dG' = RT\ln(10)\,\Delta N_{\mathrm{H}}\mathrm{dpH} \qquad \text{(Eq. S1)}$$

Therefore,

$$\Delta_{\mathrm{r}}G'^{\circ}(\mathrm{pH}) = \Delta_{\mathrm{r}}G^{\circ} + RT\ln(10)\int_{\mathrm{pH_{ref}}}^{\mathrm{pH}}\Delta_{\mathrm{r}}\overline{N_{\mathrm{H}}}\mathrm{dpH} \qquad \text{(Eq. S2)}$$

where $\Delta_{\mathrm{r}}\overline{N_{\mathrm{H}}}$ denotes the change in the average number of bound protons accompanying the reaction, which is determined by the solution pH and the p$K_a$ values of titratable groups, and where $\mathrm{pH_{ref}}$ is a constant determined as $\Delta_{\mathrm{r}}G'^{\circ}(\mathrm{pH_{ref}}) = \Delta_{\mathrm{r}}G^{\circ}$. The change in the reaction free energy between pH 6.0 and 7.3 is given by:

$$\Delta\Delta_{\mathrm{r}}G'^{\circ}(\mathrm{pH_1} \to \mathrm{pH_2}) = RT\ln(10)\int_{\mathrm{pH_1}}^{\mathrm{pH_2}}\Delta_{\mathrm{r}}\overline{N_{\mathrm{H}}}\mathrm{dpH} \qquad \text{(Eq. S3)}$$

This free-energy change can be decomposed into two contributions: a term $\Delta\Delta_{\mathrm{r}}G'^{\circ}_{\text{acyl-transfer}}(\mathrm{pH}\ 6.0\rightarrow7.3)$, arising from changes in $\Delta_{\mathrm{r}}\overline{N_{\mathrm{H}}}$ associated with the functional groups directly involved in the thioester reaction (*i.e.* the thioester bond and the nucleophile), and a second term, $\Delta\Delta_{\mathrm{r}}G'^{\circ}_{\text{other}}(\mathrm{pH}\ 6.0\rightarrow7.3)$, arising from all remaining contributions to $\Delta_{\mathrm{r}}\overline{N_{\mathrm{H}}}$. The latter may include, for example, contributions from changes in the p$K_a$ values of general acid–base catalytic residues around the thioester bond. The former can be calculated explicitly within Alberty's framework.

$$\Delta\Delta_{\mathrm{r}}G'^{\circ}(\mathrm{pH_1} \to \mathrm{pH_2}) = \Delta\Delta_{\mathrm{r}}G'^{\circ}_{\text{acyl-transfer}}(\mathrm{pH_1} \to \mathrm{pH_2}) + \Delta\Delta_{\mathrm{r}}G'^{\circ}_{\text{other}}(\mathrm{pH_1} \to \mathrm{pH_2})$$

$$\Delta\Delta_{\mathrm{r}}G'^{\circ}_{\text{acyl-transfer}}(\mathrm{pH_1} \to \mathrm{pH_2}) = RT\ln(10)\int_{\mathrm{pH_1}}^{\mathrm{pH_2}}\Delta_{\mathrm{r}}\overline{N_{\mathrm{H,\,acyl\text{-}transfer}}}\mathrm{dpH} \qquad \text{(Eq. S4)}$$

$$\Delta\Delta_{\mathrm{r}}G'^{\circ}_{\text{other}}(\mathrm{pH_1} \to \mathrm{pH_2}) = RT\ln(10)\int_{\mathrm{pH_1}}^{\mathrm{pH_2}}\Delta_{\mathrm{r}}\overline{N_{\mathrm{H,\,other}}}\mathrm{dpH}$$

#### Free-energy analysis of the thioester bond cleavage in SfbI-TED

To start with, we analyze the pH-dependent reaction free energy of intramolecular thioester bond cleavage. One plausible mechanism for the cleavage reaction is hydrolysis:

$$\mathrm{R_1\text{-}CO\text{-}S\text{-}R_2 + H_2O \rightleftharpoons R_1\text{-}COOH + R_2\text{-}SH} \qquad \text{(Eq. S5)}$$

For this cleavage reaction, $\Delta\Delta_{\mathrm{r}}G'^{\circ}_{\text{acyl-transfer}}(\mathrm{pH}\ 6.0\rightarrow7.3)$ depends on the p$K_a$ values of the carboxylic acid ($R_1$-COOH) and thiol ($R_2$-SH) groups; substituting typical solution values ($\mathrm{p}K_{\mathrm{a}}^{\mathrm{R1\text{-}COOH}} = 4.5$, $\mathrm{p}K_{\mathrm{a}}^{\mathrm{R2\text{-}SH}} = 9.1$ yields the following(*32*):

$$\Delta\Delta_{\mathrm{r}}G'^{\circ}_{\text{acyl-transfer}}(\mathrm{pH}\ 6.0 \rightarrow 7.3)$$

$$= \left[-RT\left\{\ln\left(1+10^{\mathrm{pH-pK_a^{R1-COOH}}}\right) + \ln\left(1+10^{\mathrm{pH-pK_a^{R2-SH}}}\right)\right\}\right]_{6.0}^{7.3} \qquad \text{(Eq. S6)}$$

$$= -1.77\ (\mathrm{kcal/mol})$$

The sensitivity of $\Delta\Delta_{\mathrm{r}}G'^{\circ}_{\text{acyl-transfer}}$ to variations in p$K_a$ values is provided in fig. S9.

Meanwhile, the fraction of cleaved thioester bonds at each pH was approximated from the normalized $A_{324}$ values, as a proxy for the equilibrium cleavage fraction.

$$K'(\mathrm{pH}) \approx \frac{\text{normalized } A_{324}/100}{1 - \text{normalized } A_{324}/100}$$

$$\Delta_{\mathrm{r}}G'^{\circ}(\mathrm{pH}\ 6.0) = -RT\ln(K') \approx \quad 1.59 \pm 0.12\ (\mathrm{kcal/mol}) \qquad \text{(Eq. S7)}$$

$$\Delta_{\mathrm{r}}G'^{\circ}(\mathrm{pH}\ 7.3) = -RT\ln(K') \approx -0.04 \pm 0.20\ (\mathrm{kcal/mol})$$

Here, the uncertainties represent standard deviations derived from three independent measurements. Therefore, based on these experimental values,

$$\Delta\Delta_{\mathrm{r}}G'^{\circ}(\mathrm{pH}\ 6.0{\rightarrow}7.3) = -1.63 \pm 0.24\ (\mathrm{kcal/mol}) \qquad \text{(Eq. S8)}$$

We next compare this experimental value with the theoretical value of $G'^{\circ}_{\text{acyl-transfer}}\ (\mathrm{pH}\ 6.0 \rightarrow 7.3)$.

$$\begin{aligned}
&\Delta\Delta_{\mathrm{r}}G'^{\circ}(\mathrm{pH}\ 6.0{\rightarrow}7.3) \\
&\quad = \Delta\Delta_{\mathrm{r}}G'^{\circ}_{\text{acyl-transfer}}\ (\mathrm{pH}\ 6.0 \rightarrow 7.3) + \Delta\Delta_{\mathrm{r}}G'^{\circ}_{\text{other}}(\mathrm{pH}\ 6.0{\rightarrow}7.3) \\
&-1.63 \pm 0.24 \ = -1.77 + \Delta\Delta_{\mathrm{r}}G'^{\circ}_{\text{other}}(\mathrm{pH}\ 6.0{\rightarrow}7.3) \\
&\Delta\Delta_{\mathrm{r}}G'^{\circ}_{\text{other}}(\mathrm{pH}\ 6.0{\rightarrow}7.3) = +0.14 \pm 0.24\ (\mathrm{kcal/mol})
\end{aligned} \qquad \text{(Eq. S9)}$$

This result suggests that additional pH-dependent contributions beyond the intrinsic acyl-transfer chemistry are small in the system.

**Free-energy analysis assuming aminolysis of the intramolecular thioester bond**

In the analysis above, we assumed that the cleavage detected by the thiol-quantification assay primarily reflects hydrolysis of the intramolecular thioester bond. Although no low-molecular weight amines were present in the reaction solution used for the thiol quantification experiment, it remains possible that amino groups on SfbI-TED participated in the cleavage reaction. Even in such a case, the conclusions described above remain robust, described below.

In the case in which the intramolecular thioester bond is cleaved by amino groups on SfbI-TED:

$$\mathrm{R_1\text{-}CO\text{-}S\text{-}R_2 + R_3\text{-}NH_2 \rightleftharpoons R_1\text{-}CO\text{-}NH\text{-}R_3 + R_2\text{-}SH} \qquad \text{(Eq. S10)}$$

The theoretical value of $\Delta\Delta_{\mathrm{r}}G'^{\circ}_{\text{acyl-transfer}}(\mathrm{pH})$ can be calculated as:

$$\begin{aligned}
&\Delta\Delta_{\mathrm{r}}G'^{\circ}_{\text{acyl-transfer}}(\mathrm{pH}\ 6.0{\rightarrow}7.3) \\
&\quad = -RT\left[\ln\left(1 + 10^{\mathrm{pH} - \mathrm{p}K_{\mathrm{a}}^{\text{R2-SH}}}\right) - \ln\left(1 + 10^{\mathrm{p}K_{\mathrm{a}}^{\text{R3-NH2}} - \mathrm{pH}}\right)\right]_{6.0}^{7.3}
\end{aligned} \qquad \text{(Eq. S11)}$$

Although the p$K_{\mathrm{a}}$ values for the thiol ($R_2$-SH) and amine ($R_3$-$NH_2$) groups may vary within the catalytic pocket of the TED, substituting typical solution values ($\mathrm{p}K_{\mathrm{a}}^{\text{R2-SH}} = 9.1$ and $\mathrm{p}K_{\mathrm{a}}^{\text{R3-NH2}} = 10.4$) (*32*) yields the following:

$$\Delta\Delta_{\rm r}G'^{\circ}_{\text{acyl-transfer}}(\text{pH } 6.0{\rightarrow}7.3)$$

$$= -RT\left\{\ln\left(1 + 10^{\text{pH}-\text{pK}_\text{a}^{\text{R2-SH}}}\right) - \ln\left(1 + 10^{\text{pK}_\text{a}^{\text{R3-NH2}}-\text{pH}}\right)\right\}\Bigg|_{6.0}^{7.3} \qquad \text{(Eq. S12)}$$

$$= -1.78 \text{ (kcal/mol)}$$

The sensitivity of $\Delta\Delta_{\rm r}G'^{\circ}_{\text{acyl-transfer}}$ to variations in p$K_{\rm a}$ values is the same as fig. S10.
Next, we calculate the equilibrium constant from the normalized $A_{324}$ measured in the thiol assay. Different from the equation of hydrolysis, in aminolysis the effective concentration of amino groups affects the calculated equilibrium constant. Amino groups that can participate in this reaction include N-termini and lysine side chains, either on the same molecule as the thioester bond or on a different molecule. Our construct of SfbI-TED contains 20 lysine residues and one N-terminus, including those in the tag peptide. Because the fraction of these groups that can actually participate in the reaction is unknown, we introduce a constant γ to account for the effective concentration and express the total activity of amino groups as:

$$a_{\text{R3-NH}_2\text{, total}} = \gamma \cdot a_{\text{TED}} \qquad \text{(Eq. S13)}$$

Then, the equilibrium constant can be approximated using the normalized $A_{324}$ as follows:

$$K' \approx \frac{\text{normalized } A_{324}/100}{1 - \text{normalized } A_{324}/100} \cdot \frac{1}{0.00003(\gamma - \text{normalized } A_{324}/100)} \qquad \text{(Eq. S14)}$$

From the relationship between the Gibbs free energy and the equilibrium constant,

$$\Delta\Delta_{\rm r}G'^{\circ}(\text{pH } 6.0 \rightarrow 7.3\ )$$

$$= -RT\ln\left(\frac{\text{normalized } A_{324}/100}{(1 - \text{normalized } A_{324}/100)(\gamma - \text{normalized } A_{324}/100)}\right)\Bigg|_{6.0}^{7.3} \qquad \text{(Eq. S15)}$$

This value varies as a function of γ. When the thioester bond reacts with amino groups on other molecules, γ has a maximum value of 20, whereas reactions with amino groups within the same molecule are expected to correspond to even larger effective γ values.
As an example of the value of $\Delta\Delta_{\rm r}G'^{\circ}_{\text{deacylation}}(\text{pH } 6.0{\rightarrow}7.3)$:

$$\text{For } \gamma = 1,\ -2.01 \pm 0.34 \text{ (kcal/mol)}$$
$$\text{For } \gamma = 5,\ -1.68 \pm 0.25 \text{ (kcal/mol)}$$
$$\text{For } \gamma = 20,\ -1.63 \pm 0.24 \text{ (kcal/mol)}$$

Thus, with increasing γ, the value converges to $-1.63 \pm 0.24$ kcal/mol. This asymptotic value is consistent with that calculated under the assumption of hydrolysis. As an illustrative example, when γ = 5 is adopted,

$$\Delta\Delta_{\rm r}G'^{\circ}(\text{pH } 6.0 \rightarrow 7.3)$$

$$= \Delta\Delta_{\rm r}G'^{\circ}_{\text{acyl-transfer}}(\text{pH } 6.0 \rightarrow 7.3) + \Delta\Delta_{\rm r}G'^{\circ}_{\text{other}}(\text{pH } 6.0 \rightarrow 7.3) \qquad \text{(Eq. S16)}$$

$$-1.68 = -1.78 + \Delta\Delta_{\rm r}G'^{\circ}_{\text{other}}(\text{pH } 6.0 \rightarrow 7.3)$$

$$\Delta\Delta_{\rm r}G'^{\circ}_{\text{other}}(\text{pH } 6.0 \rightarrow 7.3) = 0.1 \pm 0.25 \text{ (kcal/mol)}$$

Therefore, the same conclusion is obtained as in the case where hydrolysis is assumed.

**Free-energy analysis of the intermolecular isopeptide bond formation in the SfbI-TED–fibrinogen complex**

In the crosslinking reaction in the complex, an analogous thermodynamic analysis can be applied.

$$\mathrm{R_1\text{-}CO\text{-}S\text{-}R_2} + \mathrm{R_3\text{-}NH_2} \rightleftharpoons \mathrm{R_1\text{-}CO\text{-}NH\text{-}R_3} + \mathrm{R_2\text{-}SH} \quad \text{(Eq. S17)}$$

Although the p$K_a$ values for the thiol ($R_2$-SH) and amine ($R_3$-$NH_2$) groups may vary within the catalytic pocket of the TED, substituting typical solution values ($\mathrm{p}K_\mathrm{a}^{\mathrm{R2\text{-}SH}} = 9.1$ and $\mathrm{p}K_\mathrm{a}^{\mathrm{R3\text{-}NH2}} = 10.4$)(*32*) yields the following:

$$\Delta\Delta_\mathrm{r} G'^\circ_{\text{acyl-transfer}}(\mathrm{pH}\ 6.0 \rightarrow 7.3) = -RT \ln\left(\frac{1 + 10^{\mathrm{pH}-\mathrm{p}K_\mathrm{a}^{\mathrm{R2\text{-}SH}}}}{1 + 10^{\mathrm{p}K_\mathrm{a}^{\mathrm{R3\text{-}NH2}}-\mathrm{pH}}}\right)\Bigg|_{6.0}^{7.3} \quad \text{(Eq. S18)}$$

$$= -1.78\ (\mathrm{kcal/mol})$$

The sensitivity of $\Delta\Delta_\mathrm{r} G'^\circ_{\text{acyl-transfer}}$ to variations in p$K_a$ values is provided in fig. S10. This free-energy change refers solely to the reaction among the sidechains of cysteine, glutamine and lysine. At the macromolecular level, the overall transformed standard Gibbs energy change is given by:

$$\Delta\Delta_\mathrm{r} G'^\circ(\mathrm{pH}\ 6.0 \rightarrow 7.3\ ) = \Delta\Delta_\mathrm{r} G'^\circ_{\text{acyl-transfer}} + \Delta\Delta_\mathrm{r} G'^\circ_{\text{other}} \quad \text{(Eq. S19)}$$

In the case of intramolecular thioester bond cleavage, the contribution of $\Delta\Delta G'^\circ_{\text{other}}$ was small ($0.14 \pm 0.24$ kcal/mol) and therefore insufficient to appreciably alter the pH dependence dictated by $\Delta\Delta_r G'^\circ_{\text{acyl-transfer}}$. By the same reasoning, $\Delta\Delta_\mathrm{r} G'^\circ_{\text{other}}$ is unlikely to counterbalance the large energy shift of $\Delta\Delta_\mathrm{r} G'^\circ_{\text{acyl-transfer}} = -1.78$ kcal/mol for isopeptide bond formation. Thus, the intrinsic pH dependence of the acyl-transfer reaction is expected to manifest as the pH dependence of isopeptide bond formation.

As an order-of-magnitude estimate, assuming that p$K_a$ shifts in the complex arising from intermolecular acyl-transfer reaction are negligible (*i.e.*, $\Delta\Delta_r G'^\circ_{\text{other}} \approx 0$ kcal/mol), the transformed standard Gibbs energy at pH 6.0 can be estimated from the FASTIA-based value of $\Delta_\mathrm{r} G'^\circ(\mathrm{pH}\ 7.3) = -1.21 \pm 0.06$ (kcal/mol) (Fig. 5B), as follows:

$$\begin{aligned} \Delta_\mathrm{r} G'^\circ(\mathrm{pH}\ 6.0) &= \Delta_\mathrm{r} G'^\circ(\mathrm{pH}\ 7.3) - \Delta\Delta_\mathrm{r} G'^\circ(\mathrm{pH}\ 6.0 \rightarrow 7.3) \\ &= -1.21 - (-1.78 - 0)(\mathrm{kcal/mol}) \\ &= +\ 0.57\ (\mathrm{kcal/mol}) \end{aligned} \quad \text{(Eq. S20)}$$

**pH responsiveness of the thioester-bond cleavage in CpTIE-TED**

Assuming that cleavage of the intramolecular thioester bond in CpTIE-TED proceeds via either hydrolysis or aminolysis by a large excess of amine, the reaction free energy of the cleavage reaction at each pH can be expressed based on the thiol-quantification measurements as follows:

$$\begin{aligned} \Delta_\mathrm{r} G'^\circ(\mathrm{pH}\ 6.0) &= -RT \ln(K') \approx \ \ 1.02 \pm 0.15\ (\mathrm{kcal/mol}) \\ \Delta_\mathrm{r} G'^\circ(\mathrm{pH}\ 7.3) &= -RT \ln(K') \approx -0.26 \pm 0.05\ (\mathrm{kcal/mol}) \end{aligned} \quad \text{(Eq. S21)}$$

Therefore,

$$\Delta\Delta_r G'^\circ(\text{pH } 6.0 \rightarrow 7.3) = \Delta_r G'^\circ(\text{pH } 7.3) - \Delta_r G'^\circ(\text{pH } 6.0)$$
$$= -1.28 \pm 0.16 \text{ (kcal/mol)} \qquad \text{(Eq. S22)}$$

Comparing this experimental value with the theoretical value of $\Delta\Delta_r G'^\circ_{\text{acyl-transfer}}\,(\text{pH } 6.0 \rightarrow 7.3)$.

$$\Delta\Delta_r G'^\circ(\text{pH } 6.0{\rightarrow}7.3)$$
$$= \Delta\Delta_r G'^\circ_{\text{acyl-transfer}}\,(\text{pH } 6.0 \rightarrow 7.3) + \Delta\Delta_r G'^\circ_{\text{other}}(\text{pH } 6.0{\rightarrow}7.3) \qquad \text{(Eq. S23)}$$
$$-1.28 \pm 0.16 = -1.77 + \Delta\Delta_r G'^\circ_{\text{other}}(\text{pH } 6.0{\rightarrow}7.3)$$
$$\Delta\Delta_r G'^\circ_{\text{other}}(\text{pH } 6.0{\rightarrow}7.3) = 0.49 \pm 0.16 \text{ (kcal/mol)}$$

This indicates that the pH responsiveness of the acyl-transfer reaction becomes manifested at the macromolecular level of CpTIE-TED.

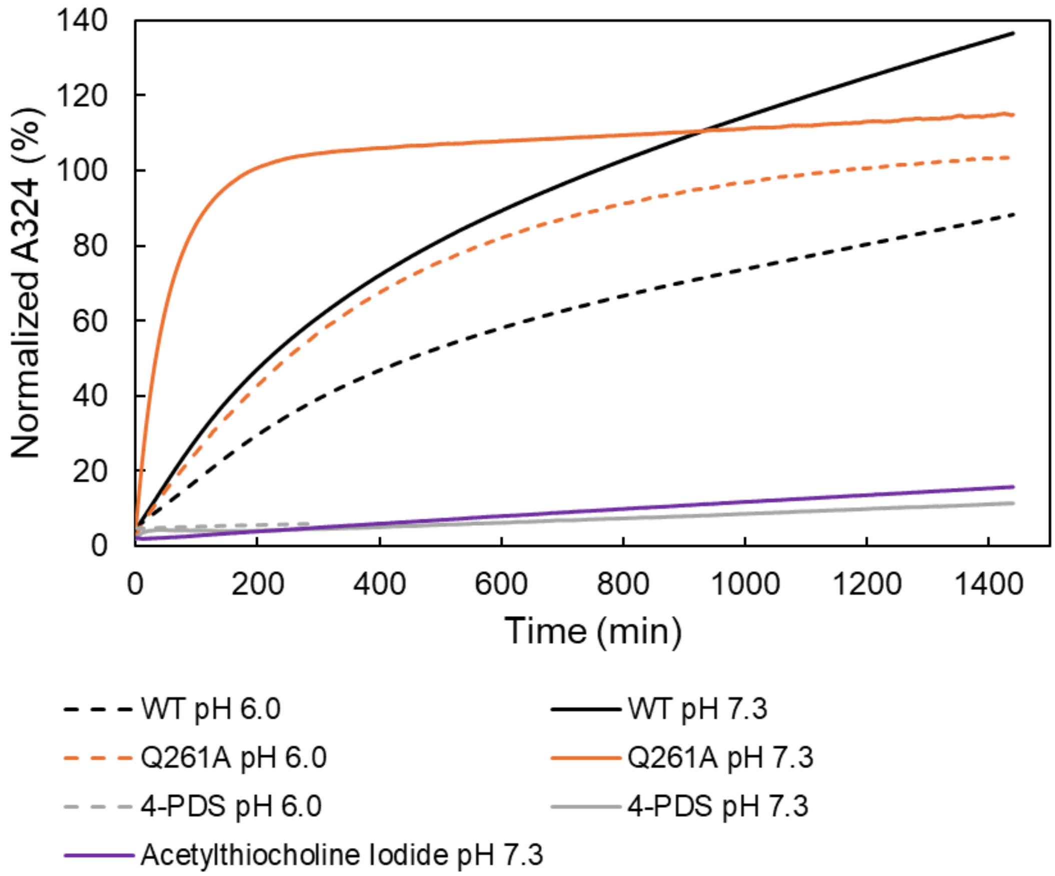


**Fig. S1. The reaction of folded SfbI-TED with 4-PDS**

Reaction between 30 µM SfbI-TED and 100 µM 4-PDS at room temperature. The vertical axis (Normalized $A_{324}$, %) was calculated by dividing the observed $A_{324}$ by the value expected for complete thioester-bond cleavage in 30 µM SfbI-TED. Solid lines indicate reactions at pH 7.3, and dashed lines indicate reactions at pH 6.0. The Q261A mutant (orange) contains one free thiol. In the wild type (black), a single thiol is generated only upon cleavage of the intramolecular thioester bond. Acetylthiocholine iodide was monitored as a model compound for a small-molecule thioester bond.

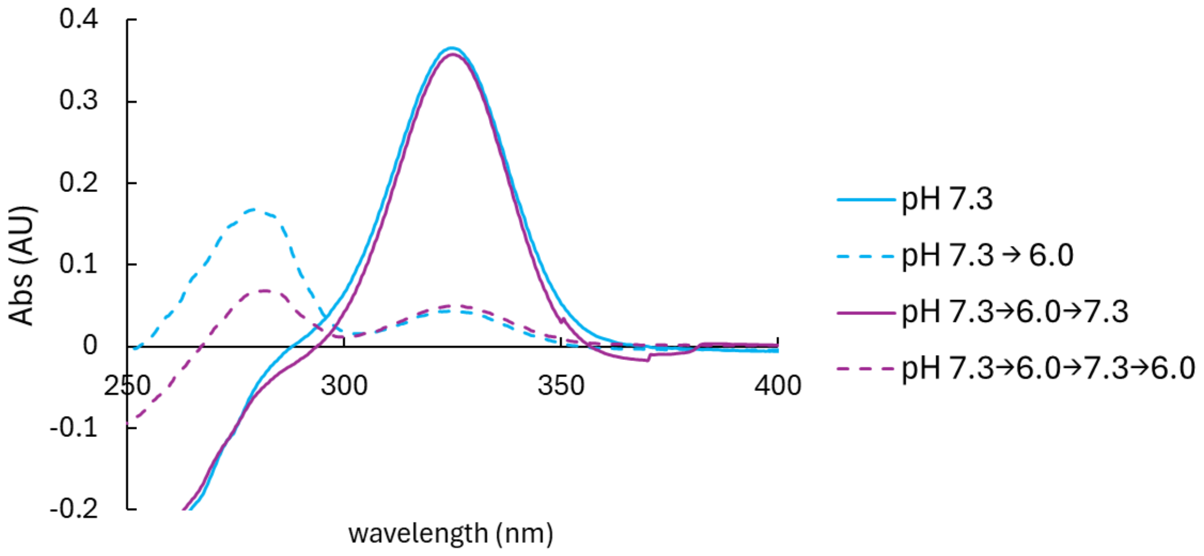


**Fig. S2. The retained pH responsiveness after repeated pH cycle**

Reaction between 30 µM SfbI-TED and 100 µM 4-PDS in solutions whose pH was exchanged by repeated dialysis. The absorbance at 324 nm was determined by the pH at the time of measurement, independent of the dialysis history.

**A.**

WT : S E N E D Y Q N L L S A E Y V P

Q261E conversion: S E N E D Y E N L L S A E Y V P

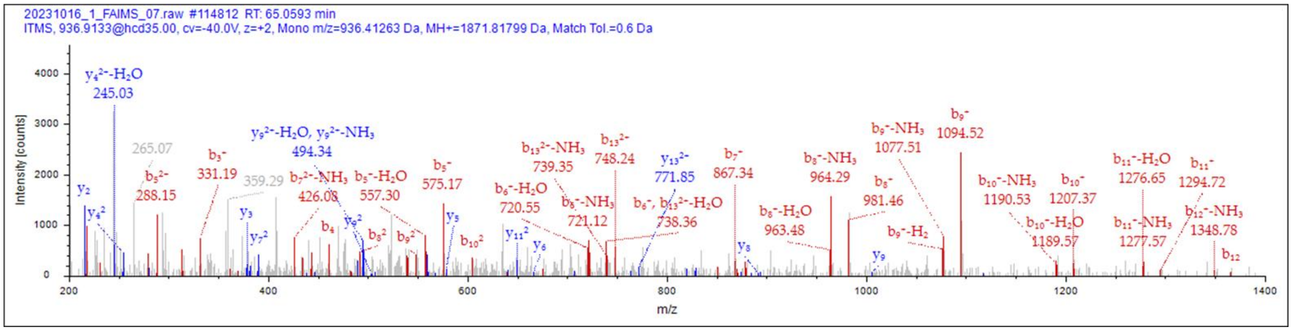


**B.**

| #1 | b⁺ | b²⁺ | Seq. | y⁺ | y²⁺ | #2 |
|---|---|---|---|---|---|---|
| 1 | 88.03930 | 44.52329 | S | | | 16 |
| 2 | 217.08190 | 109.04459 | E | 1784.78612 | 892.89670 | 15 |
| 3 | 331.12483 | 166.06605 | N | 1655.74352 | 828.37540 | 14 |
| 4 | 460.16742 | 230.58735 | E | 1541.70060 | 771.35394 | 13 |
| 5 | 575.19436 | 288.10082 | D | 1412.65800 | 706.83264 | 12 |
| 6 | 738.25769 | 369.63248 | Y | 1297.63106 | 649.31917 | 11 |
| 7 | 867.30028 | 434.15378 | E | 1134.56773 | 567.78750 | 10 |
| 8 | 981.34321 | 491.17524 | N | 1005.52514 | 503.26621 | 9 |
| 9 | 1094.42727 | 547.71728 | L | 891.48221 | 446.24474 | 8 |
| 10 | 1207.51134 | 604.25931 | L | 778.39815 | 389.70271 | 7 |
| 11 | 1294.54337 | 647.77532 | S | 665.31408 | 333.16068 | 6 |
| 12 | 1365.58048 | 683.29388 | A | 578.28205 | 289.64467 | 5 |
| 13 | 1494.62307 | 747.81517 | E | 507.24494 | 254.12611 | 4 |
| 14 | 1657.68640 | 829.34684 | Y | 378.20235 | 189.60481 | 3 |
| 15 | 1756.75482 | 878.88105 | V | 215.13902 | 108.07315 | 2 |
| 16 | | | P | 116.07061 | 58.53894 | 1 |

**Fig. S3. MS/MS analysis of the peptide with a Q261E conversion arising from hydrolysis of the thioester bond**

MS/MS spectrum and fragment ion assignment of the peptide SENEDYENLLSAEYVP. A. Detected b- and y-type fragment ions are annotated in red and blue, respectively. B. Table of the corresponding fragment ions assigned to the theoretical m/z values, confirming sequence coverage across the entire peptide.

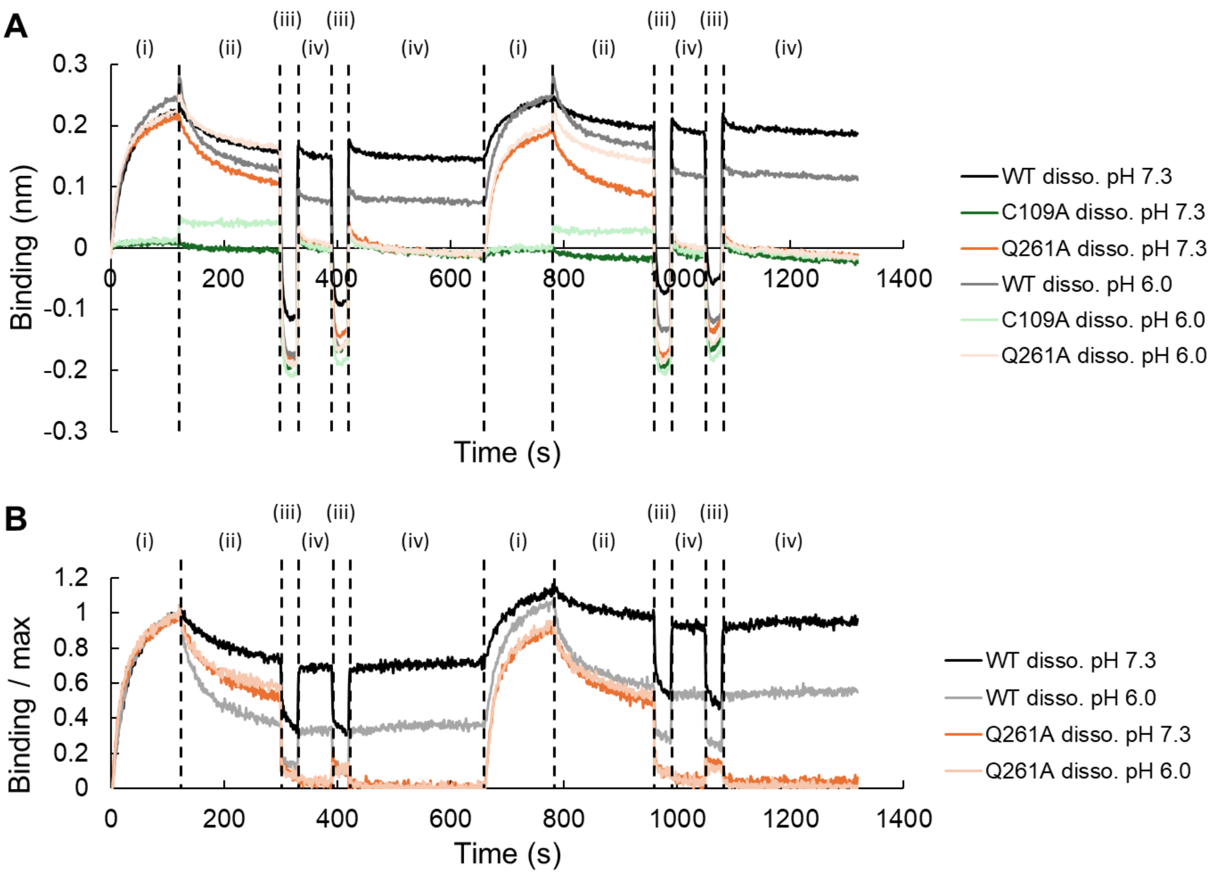


**Fig. S4. Interaction of SfbI-TED WT and mutants with fibrinogen**

BLI sensorgrams display the sequential kinetic steps consisting of (i) association with fibrinogen in pH 7.3, (ii) dissociation in either pH 7.3 or pH 6.0, (iii) strong wash in pH 2.0, and (iv) neutralization in pH 7.3. A. Raw data. B. Processed data. Sensorgrams of the SfbI-TED C109A mutant were used as a reference and subtracted to remove noise arising from nonspecific adsorption of the analyte and buffer changes. The vertical axis represents the response normalized to the maximum value in the association step.

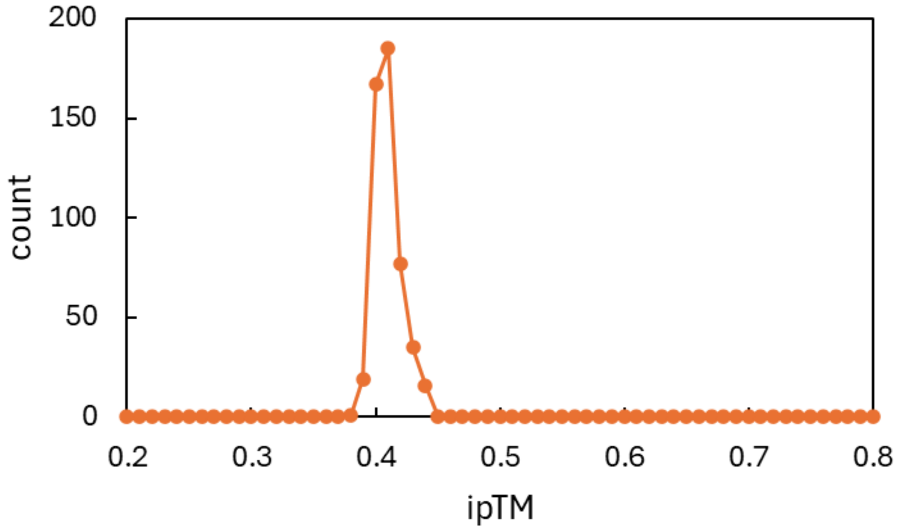


**Fig. S5. Histogram of ipTM scores from AF3 predictions**

ipTM scores obtained from AlphaFold 3–based complex structure prediction between SfbI-TED WT and the coiled-coil region of the fibrinogen trimer (Aα, Bβ, and γ chains). A total of 500 structures were generated using 100 random seeds.

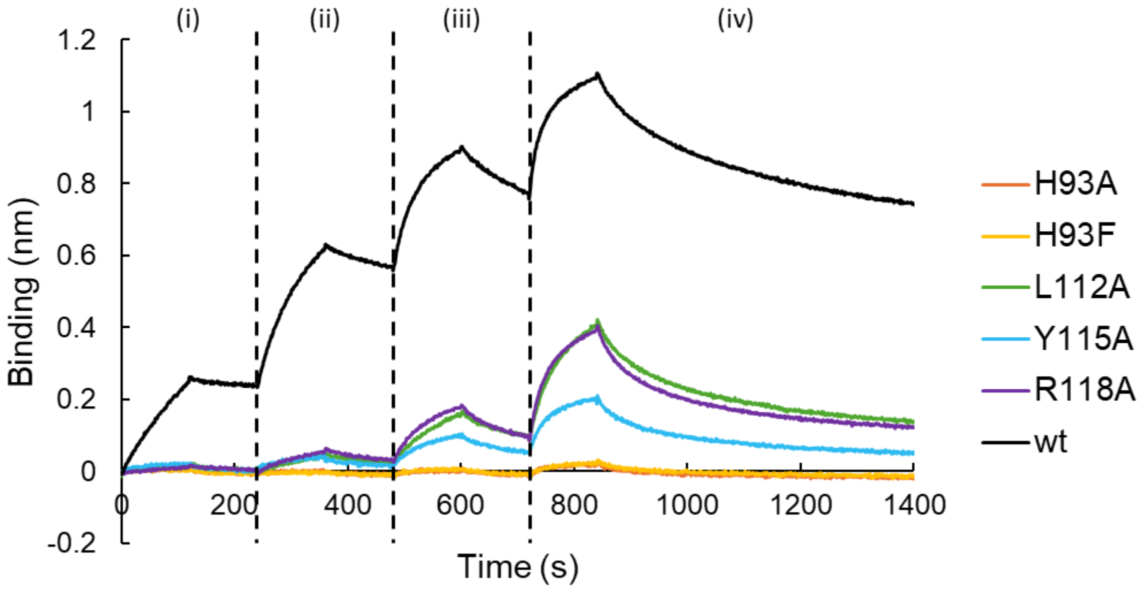


**Fig. S6. Raw BLI sensorgrams of mutant analyses performed using FASTIA**

The sensorgrams display the kinetic sequence consisting of (i)-(iv) association and dissociation steps at pH 7.3 across fibrinogen concentrations of 25, 100, 400, and 1600 nM, respectively. The H93F mutant was used as a reference for background subtraction in subsequent analyses.

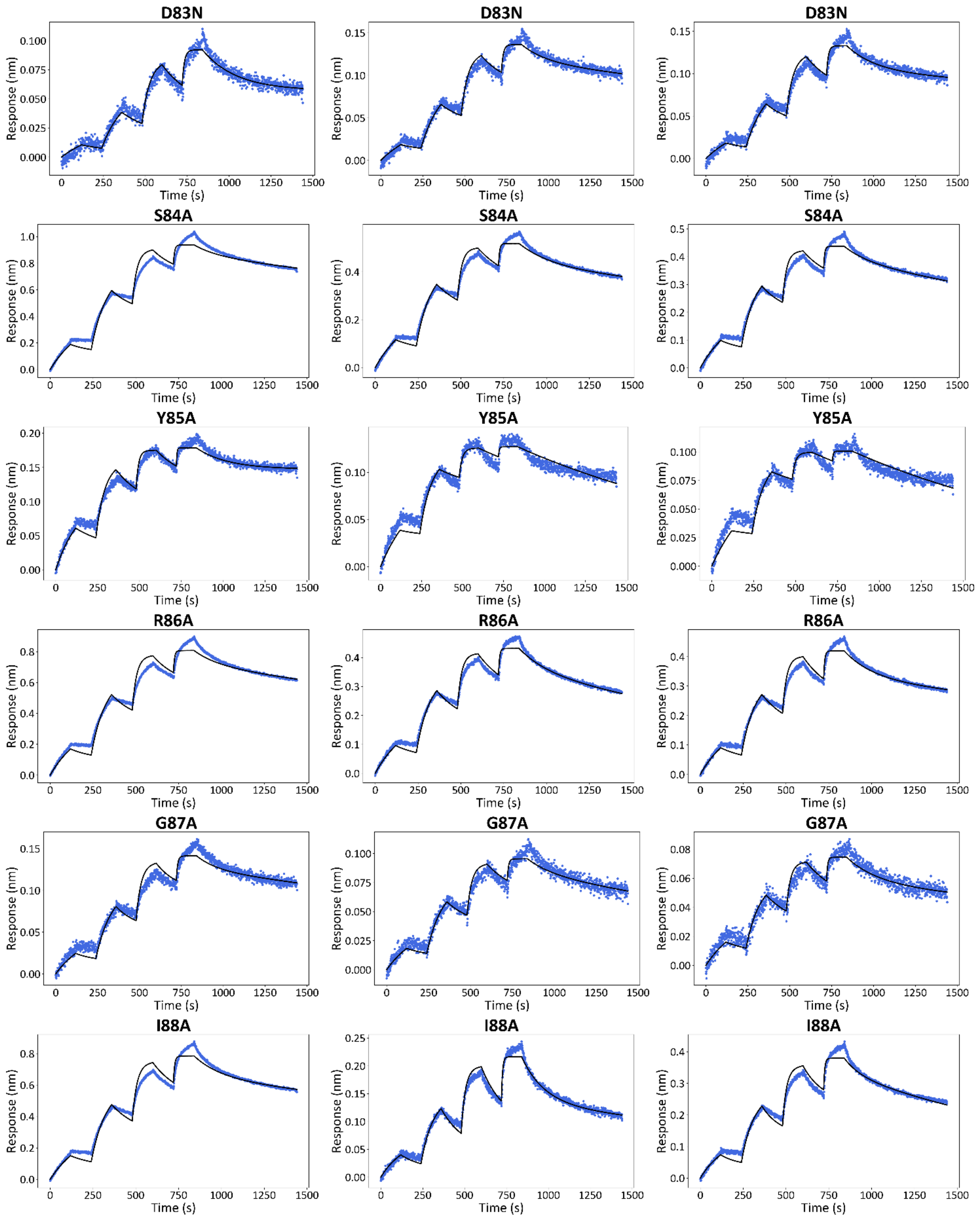


**Fig. S7. Two-step model fitting of BLI sensorgrams from FASTIA-based mutant analyses.** Blue dots represent the experimental data after subtraction of the H93F reference signal, and black solid lines indicate the fitted curves.

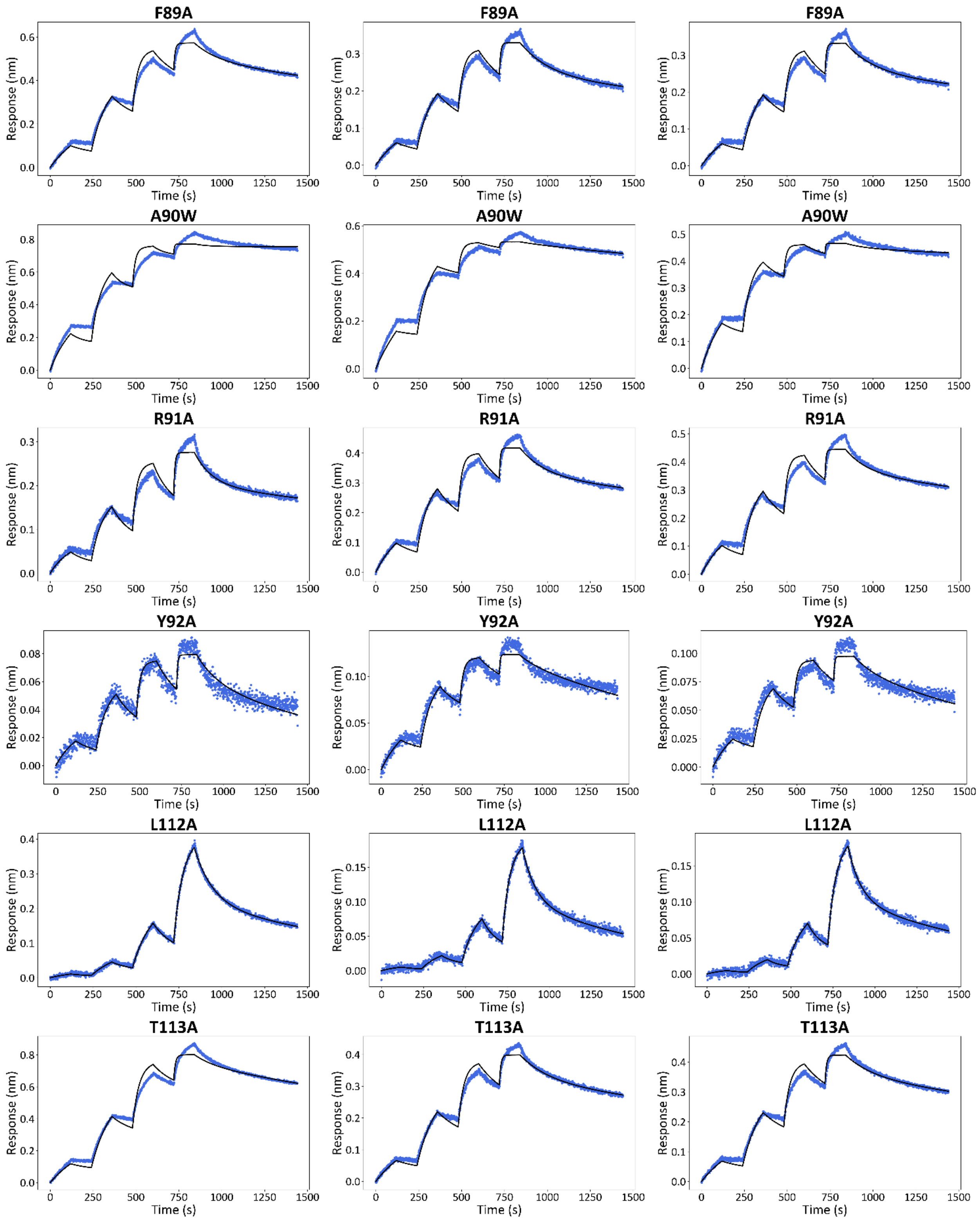


**Fig. S7  (continued)**

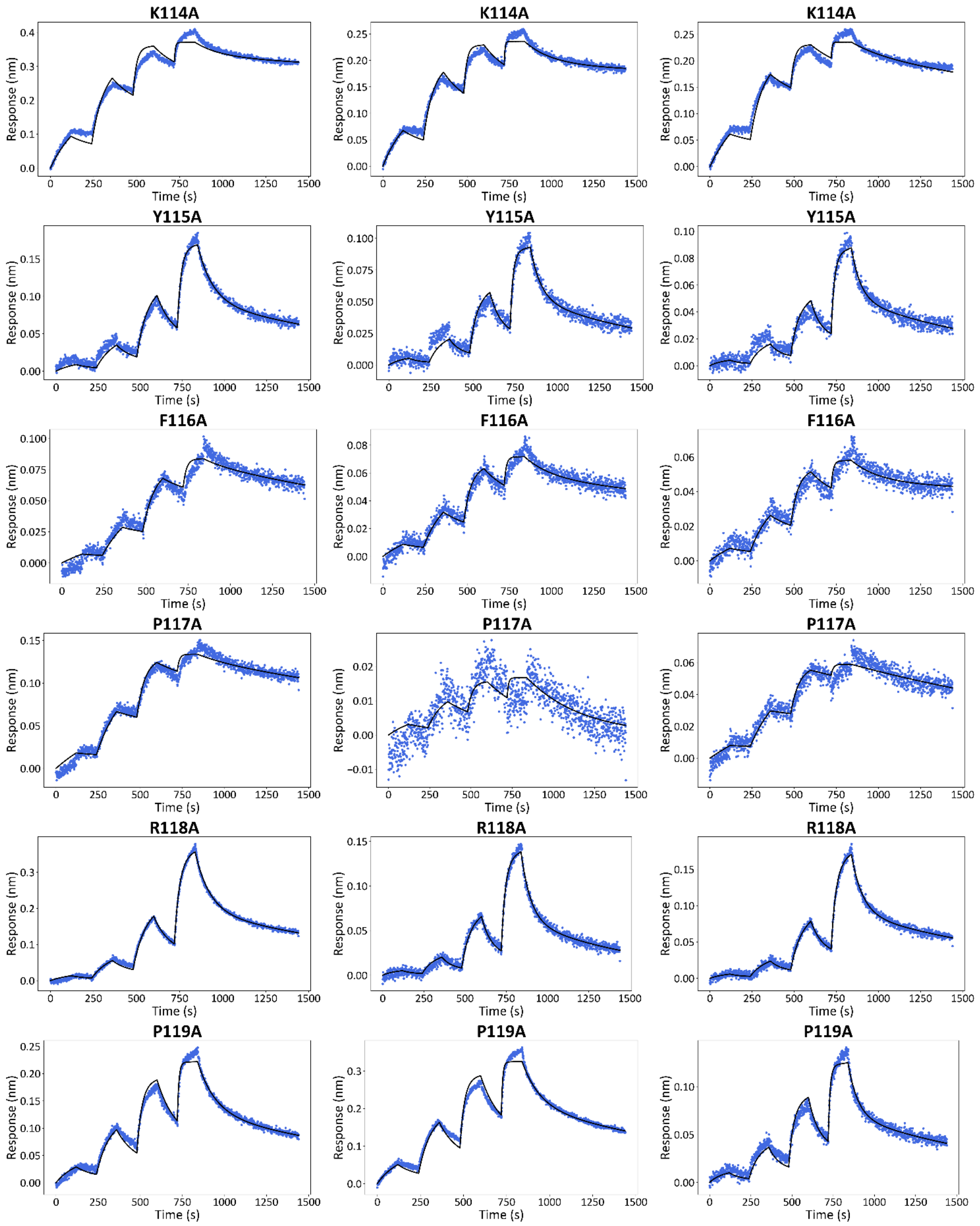


**Fig. S7 (continued)**

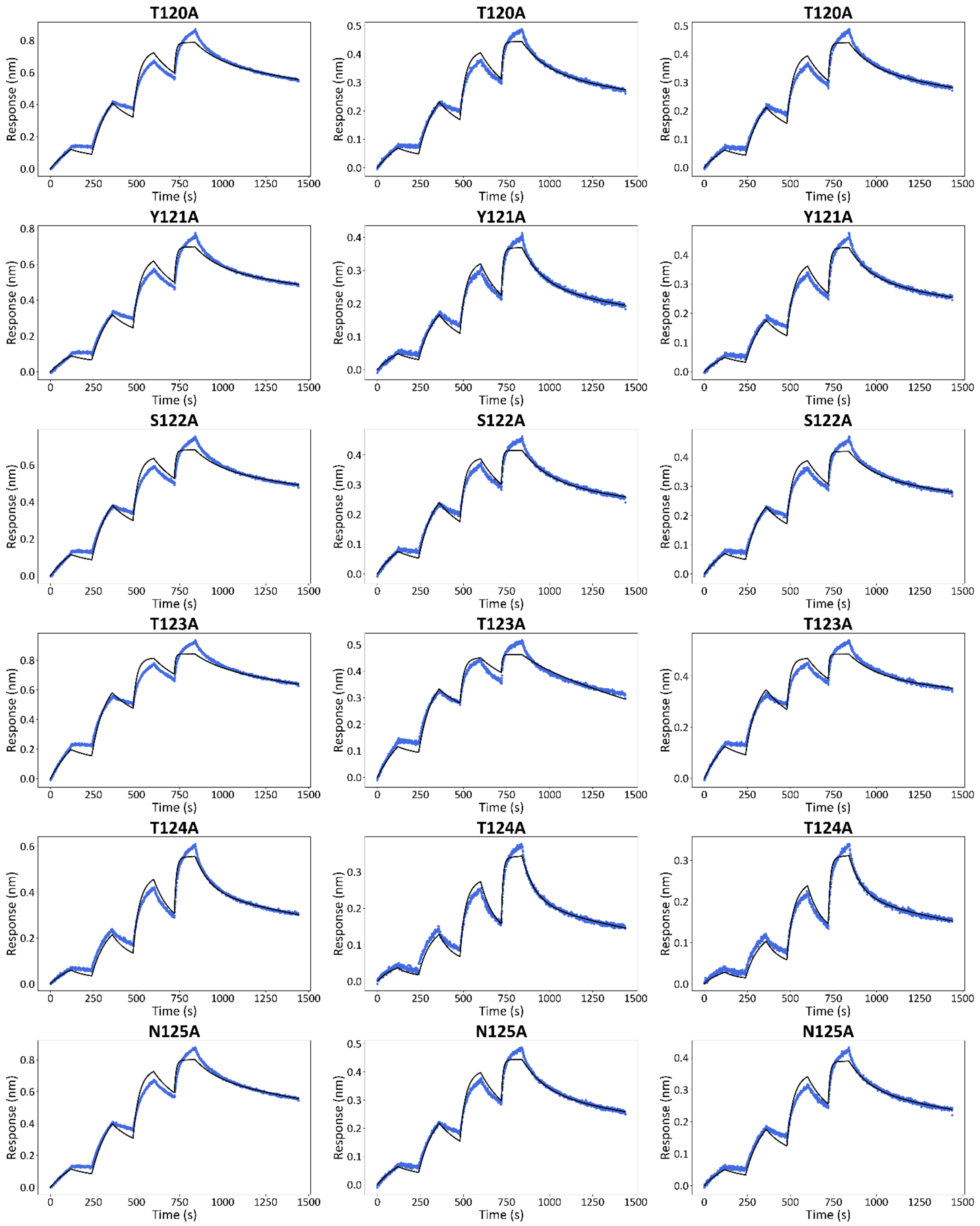


**Fig. S7  (continued)**

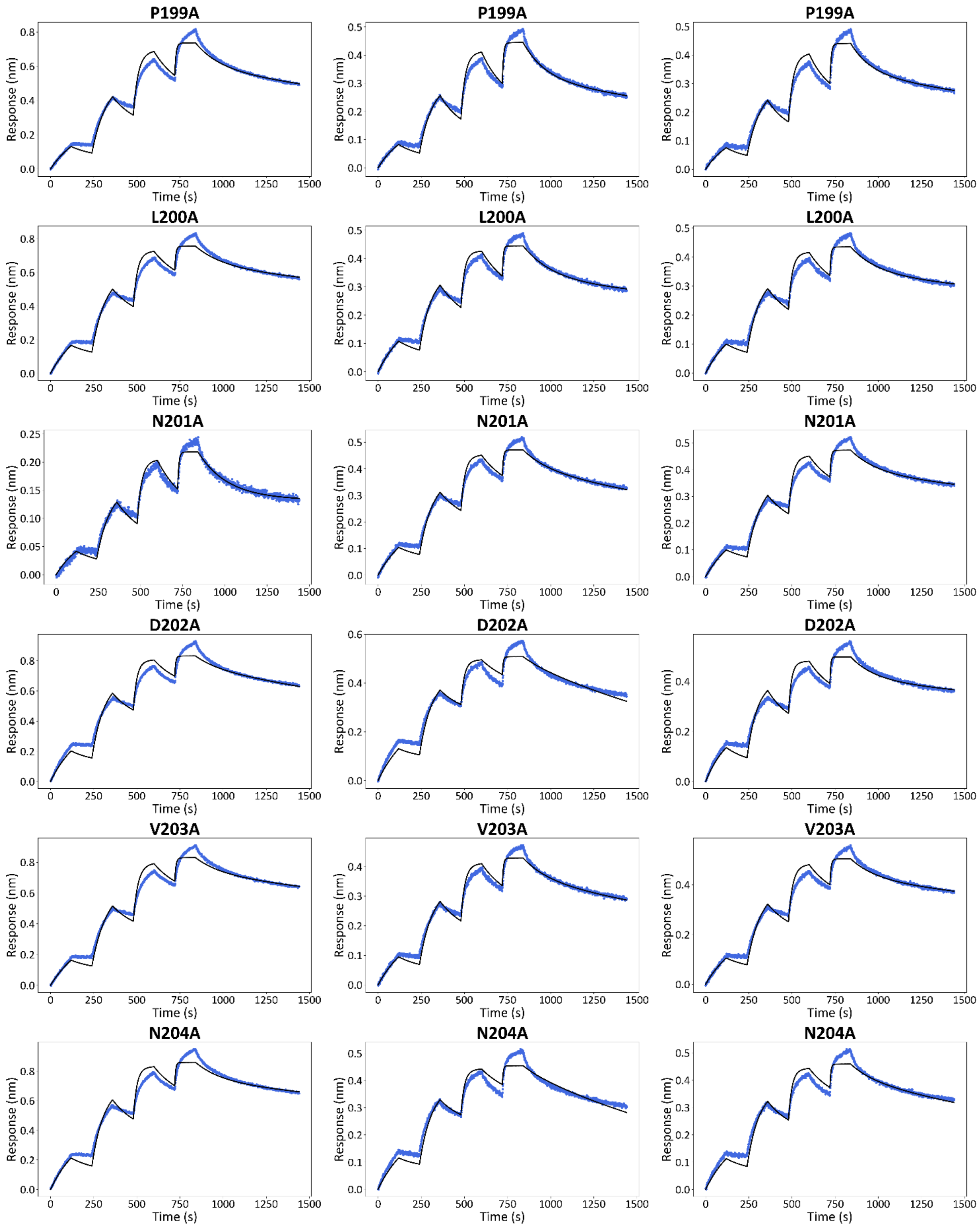


**Fig. S7 (continued)**

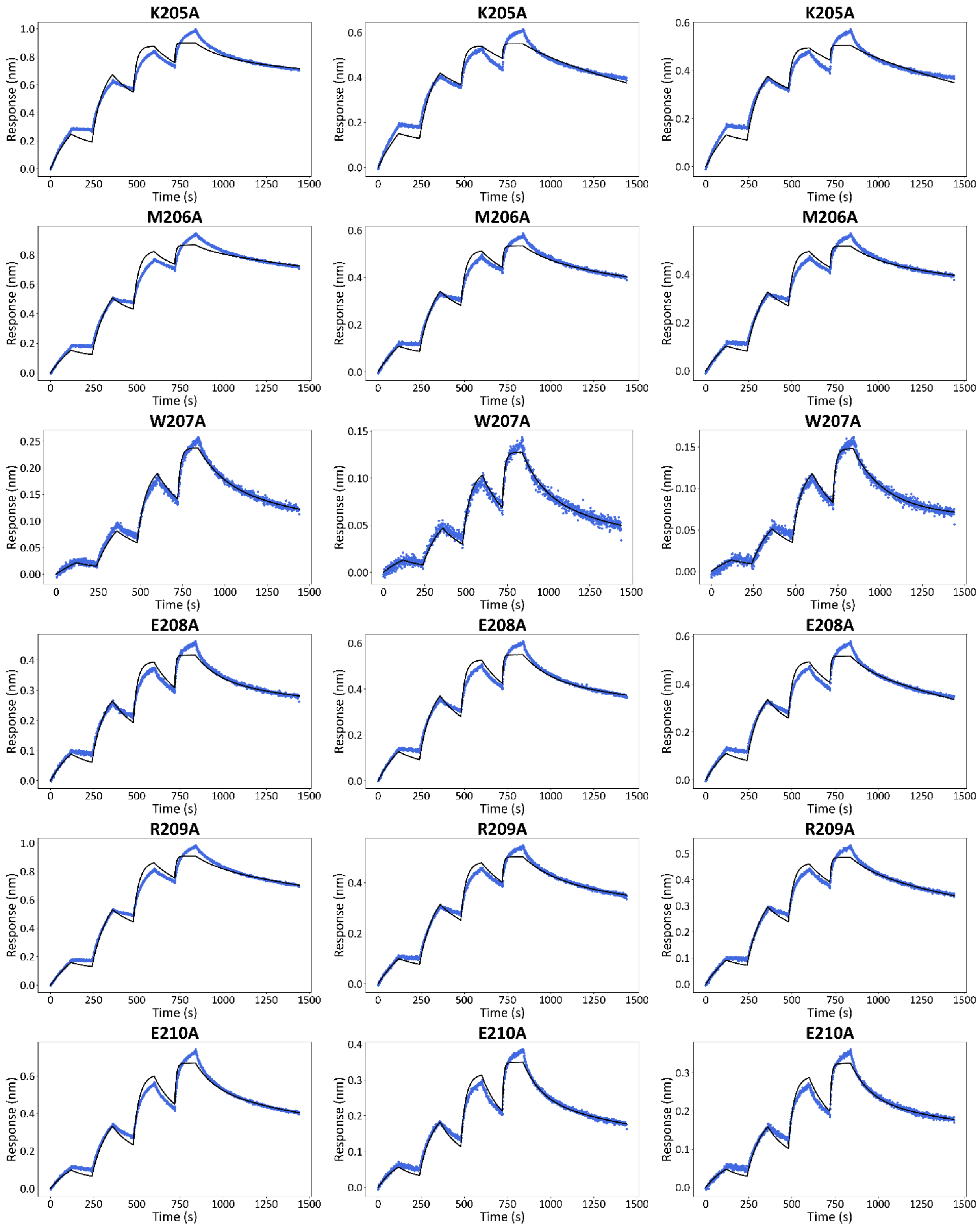


**Fig. S7 (continued)**

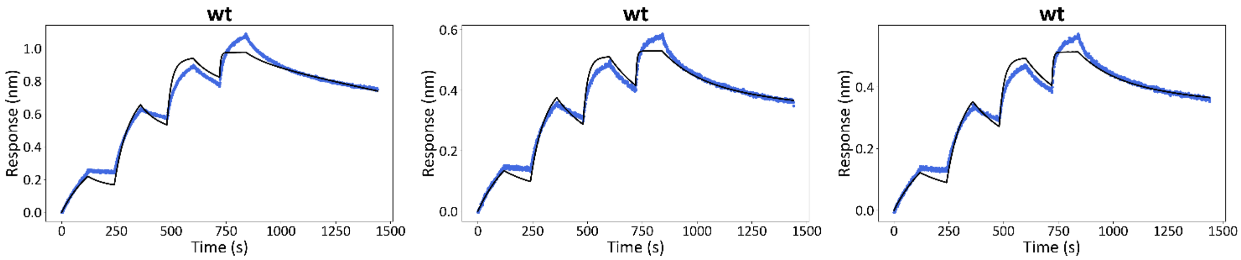


**Fig. S7 (continued)**

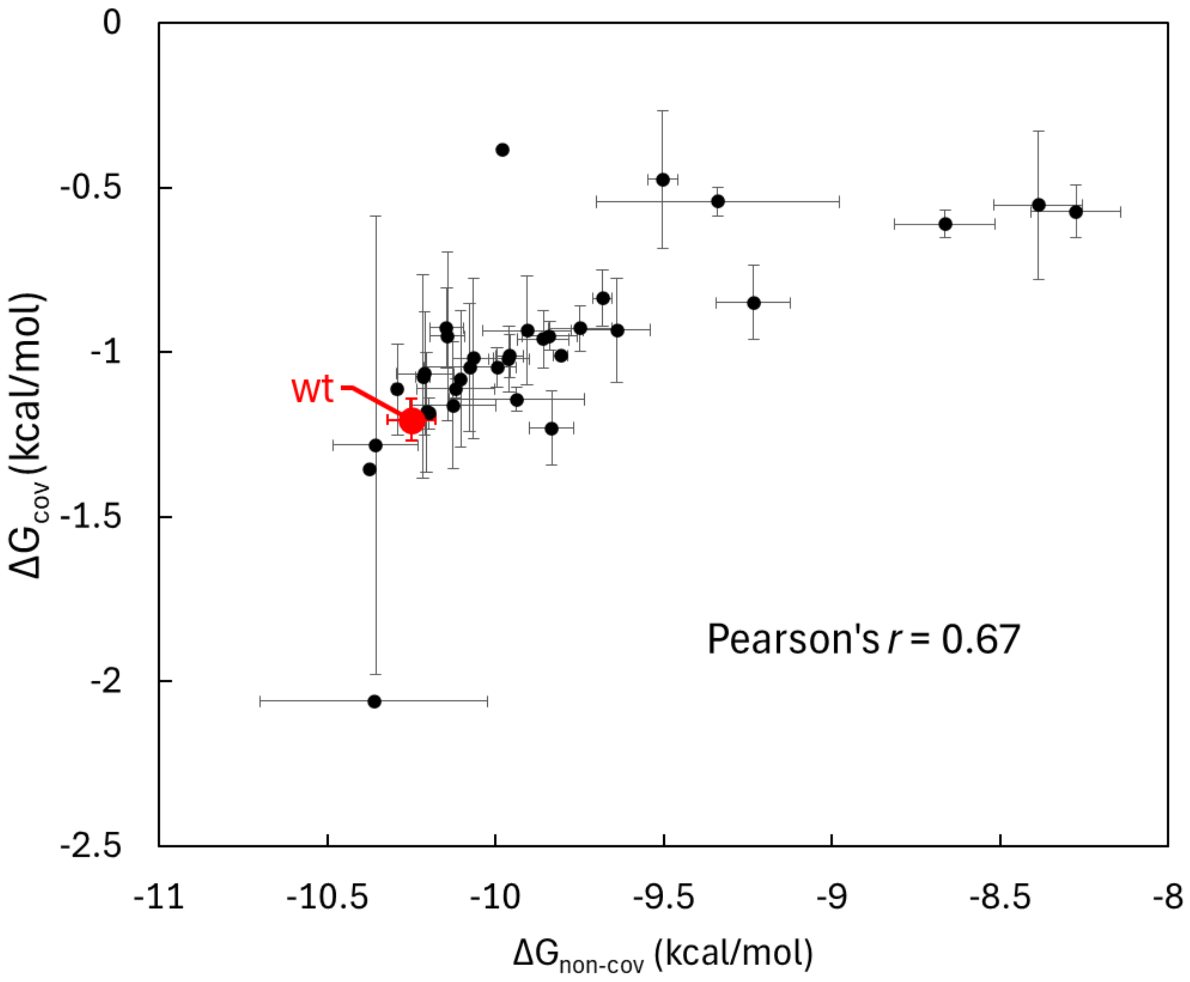


**Fig. S8. Error representation in equilibrium constant scatter plots**

Reaction free energies derived from kinetic parameters ($k_{on}$, $k_{off}$, $k_c$, and $k_{c,rev}$) obtained by two-step model fitting of BLI sensorgrams from FASTIA-based mutant analyses. The free energy of non-covalent molecular recognition (x-axis) is plotted against that of intermolecular crosslinking (y-axis). Among triplicate measurements, only data satisfying the fitting reliability criteria described in the main text were used for statistical analysis. Points represent mean values, and error bars indicate standard deviations. Error bars are not shown for mutants with only a single reliable dataset.

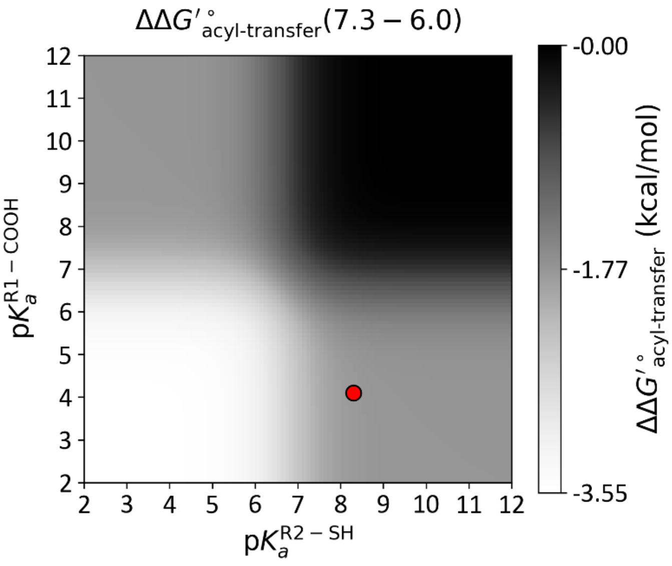


**Fig. S9. pKa-dependent changes in the reactivity of thioester bond hydrolysis**

Red dots indicate the reference values reported in the discussion. When the pKa values of the reactive groups are sufficiently distant from the pH values examined (6.0 and 7.3), the $\Delta\Delta_{\mathrm{r}}G'^{\circ}_{\text{acyl-transfer}}$ values remain robust.

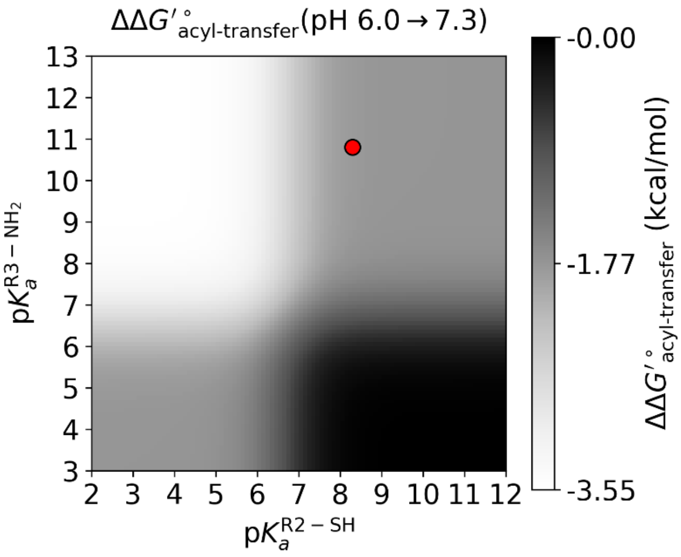


**Fig. S10. pKa-dependent changes in the reactivity of acyl transfer**

Red dots indicate the reference values reported in the discussion. When the pKa values of the reactive groups are sufficiently distant from the pH values examined (6.0 and 7.3), the $\Delta\Delta_{\mathrm{r}}G'^{\circ}_{\text{acyl-transfer}}$ values remain robust.

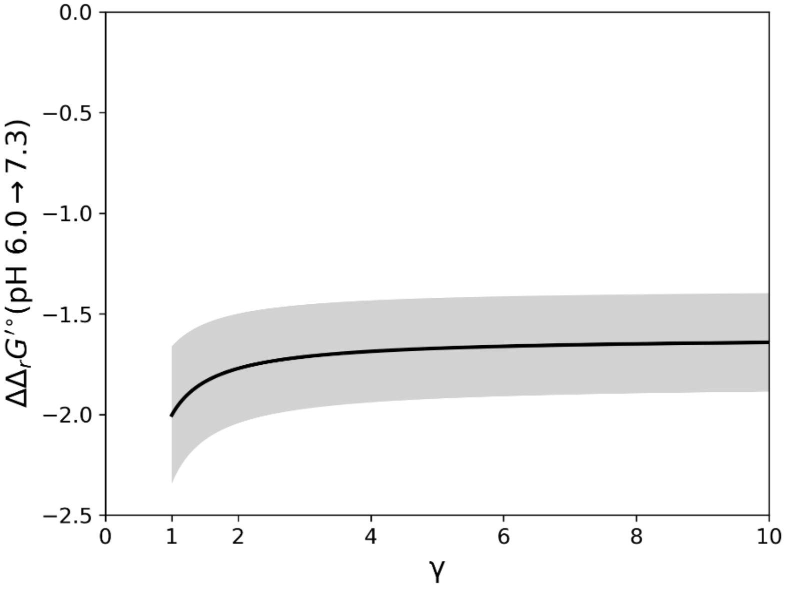


**Fig. S11. Dependence of ΔΔ$_r$*G*'°(pH 6.0 → 7.3) on γ**

The black line represents the mean value derived from three independent thiol quantification measurements, and the gray shaded area indicates the propagated uncertainty corresponding to the standard deviation.

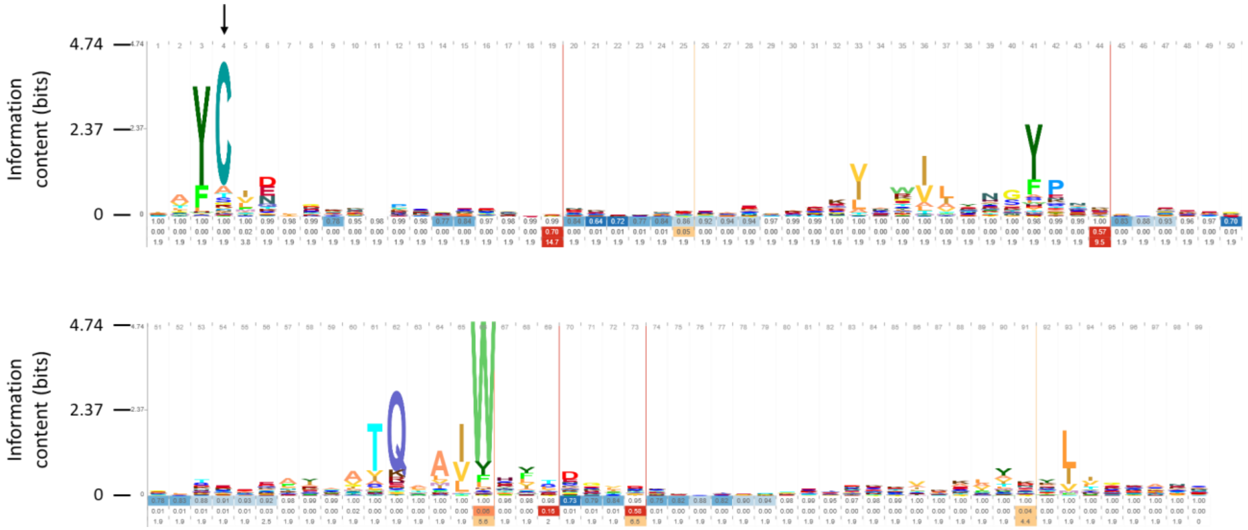


**Fig. S12. Residue conservation inferred from the Pfam TED domain profile HMM**

Sequence conservation was visualized as information content (bits) derived directly from the Pfam TED domain (PF08341) profile hidden Markov model, without reliance on sequence sampling or multiple sequence alignment of individual TED sequences. An arrow indicates the cysteine residue that forms the thioester bond.

**Table S1. DNA sequences**

| protein | DNA sequence |
|---|---|
| SfbI-TED | ATGAATCACAAAGTGCATATGGGCAGCAGCCATCATCATCATCATCACCGCGGCGTG<br>CCGCATATTGTGATGGTGGATGCGTATAAACGCTATAAAGACGAAAAACCGTCCCT<br>AACTTCAAATCCCCAGATCCTGACTATCCATGGTACGGGTACGATTCCTATCGTGGTA<br>TTTTTGCACGCTACCATAATCTGAAGGTTAATCTTAAAGGTTCGAAGGAATACCAGGC<br>GTATTGTTTCAATTTAACAAAGTACTTCCCGCGTCCCACTTACTCAACCACGAACAAC<br>TTCTACAAAAAGATTGACGGGTCCGGCAGTGCGTTCAAAAGCTACGCCGCTAATCC<br>CCGTGTATTAGACGAAAATTTAGACAAACTGGAAAAAAACATCTTAAATGTTATCTACA<br>ATGGCTACAAGAGTAACGCTAACGGGTTCATGAACGGAATCGAGGACTTAAACGCTA<br>TCTTAGTAACACAAAATGCGATTTGGTATTACTCAGATTCAGCGCCCTTGAACGATGT<br>AAATAAAATGTGGGAGCGCGAGGTCCGTAATGGTGAGATCTCAGAGTCACAAGTTA<br>CACTGATGCGTGAGGCGCTTAAAAAACTTATCGACCCGAACTTAGAAGCTACTGCTG<br>CGAATAAAATCCCTTCTGGTTACCGTCTTAATATCTTCAAATCGGAAAATGAAGATTAT<br>CAGAATTTGCTGTCCGCGGAATACGTGCCATAG |
| FbaB-TED | ATGAATCACAAAGTGGAAACTCGTAATGGAGCAAATAAACAAGGCGCTTTTGAAATC<br>AAAAAGAATAAATCCCAAGAAGAGTACAACTACGAAGTGTACGACAACCGCAACATT<br>TTACAGGACGGTGAGCACAAGTTGGAAATCAAACGTGTAGACGGCACCGGGAAAA<br>CGTACCAAGGCTTCTGTTTCCAGTTAACGAAAAACTTTCCCACGGCTCAGGGTGTTA<br>GCAAGAAGCTGTATAAAAAACTTTCTTCCTCAGATGAGGAGACACTTAAGCAGTATG<br>CAAGCAAGTACACTTCAAACCGTCGCGGTGATACAAGTGGAAATCTTAAAAAGCAAA<br>TCGCGAAGGTACTTACCGAAGGCTATCCAACGAATAAGAGTGACTGGCTGAACGGG<br>CTGACTGAGAACGAGAAAATCGAAGTCACGCAGGATGCCATTTGGTACTTTACCGA<br>AACCACAGTTCCTGCGGATCGTTCATATACGAATCGCAACGTCAATTCTCAGAAGAT<br>GAAGGAAGTGTATCAGAAGTTAATCGATACTACCGACATCGACAAGTATGAAGACGT<br>TCAATTTGACCTGTTTGTCCCCCAGGACACGAATTTACAAGCTGTTATTTCCGTAGA<br>GCCCGTCATCGAATCTTTGCCGTGGACCAGTTTGAAACCTATTGCTCAGAAAGATAT<br>TACAGCCAAAAAAATCTGGGTGGATGCACCCAAAGAAAAGCCTATTATCTACTTTAAG<br>CTGTATCGCCAGCTTCCCGGCGAAAAAGAAGTCGCAGTAGATGACGCGGAATTGAA<br>ACAAATTAATTCAGAAGGGCAACAGGAGATTAGCGTTACGTGGACTAATCAATTGGT<br>TACGGATGAGAAAGGAATGGCTTATATTTATTCGGTTAAGGAAGTGGATAAAAACGGC<br>GAGCTGTTAGAGCCTAAGGATTACATTAAAAAAGAAGATGGACTGACCGTGACCAAC<br>ACCTATGTCAAGCCTCATCATCATCATCATCACCGCGGCGTGCCGCATATTGTGATG<br>GTGGATGCGTATAAACGCTATAAATAG |

**Table S1 (continued)**

| protein | DNA sequence |
|---|---|
| CpTIE-TED | ATGAATCACAAAGTGCATATGGGCAGCAGCCATCATCATCATCATCACGGCAGCGGC<br>CTGGTGCCGCGCGGCAGCGCTAGCATGTCGGACTCAGAAGTCAATCAAGAAGCTA<br>AGCCAGAGGTCAAGCCAGAAGTCAAGCCTGAGACTCACATCAATTTAAAGGTGTCC<br>GATGGATCTTCAGAGATCTTCTTCAAGATCAAAAAGACCACTCCTTTAAGAAGGCTG<br>ATGGAAGCGTTCGCTAAAAGACAGGGTAAGGAAATGGACTCCTTAAGATTCTTGTAC<br>GACGGTATTAGAATTCAAGCTGATCAGACCCCTGAAGATTTGGACATGGAGGATAAC<br>GATATTATTGAGGCTCACAGAGAACAGATTGGTGGTATGAATTTTGGGTCGCACTTC<br>GTCTTTGCGTCAGAGAACCCCAAGATCCTGATTAAATTCTCGAATACGGAAAGTAAC<br>ACAAAGACCGAATTAAAGGGAGCAAGTTTCAAAATTGTCAAAGGGACGGACCCATC<br>CGGTCCGCCGGTAGACGGACTTAGTTGGGTCTCAGATGGCAAAATTAAGGAATTTA<br>AACTTGAGTCCGGAACGTACACTCTTGTTCAAGTCTCCGTACCAAAAGGTTACATCA<br>AAGCGGACCCTATTACATTTACCGTTAGCCCCACTGGTGGGCTGCAAACATCGACG<br>AAGTACAAGGGTTACACGTTGCTGGACAAATACCCCAAGGAAGATGATTTTCGCGAT<br>GCCATTTACATTGAGGATATGGACAATAATGACACCAGTTCCGTGGTTTATTGCTTCA<br>ATGTAACTAAGGCCACACCTACGTTCAAAGGCTCTGTGGTTAAAGTGTTATACAATGA<br>ACAATTCGGAAGCTCGAAGTTATTTACAGAGAAAGCGATCAAACCACGTGTCAAAGG<br>TGATGAACTGAAAAATAGTGTGCTTCGTGTGATCTACAACGGTTATCCCTCAAACGC<br>ATTAGGTATTAAAGAAAAATATCAACTTACAGAGGGGCAGTTCCGTAAACTTACACAG<br>CGTGCCGTGTGGAATTTTACCGATTCTAATCTTTCACTGGATAAACTGAGCCAGAAA<br>GAGATTGACGCATTAAATGAACTGATCAATGCTAAAAACGCTATCCCAGATAACCTGG<br>TCCTTAACCTTTATCTGCCCGATGACTCTTATTACCAAAACTTACTGGGTACGAAGTT<br>CGTGACTCCCAACTTAATCAAGTTAGAAAACGAGAAACTCCCCTAG |
| His-SpyCatcher-Avi | ATGAATCACAAAGTGCATCATCATCATCATCATGTTACCACTCTTTCCGGTTTGTCGG<br>GCGAACAAGGCCCAAGCGGTGATATGACAACTGAAGAAGACTCTGCTACACATATTA<br>AATTTTCCAAGCGCGACGAAGATGGCCGTGAACTGGCAGGTGCAACAATGGAATTA<br>CGTGACTCCTCCGGGAAGACAATTAGCACATGGATCAGCGACGGCCATGTAAAAGA<br>TTTCTATTTATATCCTGGAAAATACACATTCGTCGAGACTGCCGCCCCAGACGGTTAT<br>GAGGTAGCGACTCCGATTGAGTTTACCGTTAATGAAGACGGACAAGTAACCGTTGAT<br>GGAGAAGCCACCGAGGGCGATGCCCACACGGGCCTGAACGATATCTTCGAAGCGC<br>AGAAAATTGAATGGCATGAATAG |
| His-SpyCatcher | ATGAATCACAAAGTGCATCATCATCATCATCATGTTACCACTCTTTCCGGTTTGTCGG<br>GCGAACAAGGCCCAAGCGGTGATATGACAACTGAAGAAGACTCTGCTACACATATTA<br>AATTTTCCAAGCGCGACGAAGATGGCCGTGAACTGGCAGGTGCAACAATGGAATTA<br>CGTGACTCCTCCGGGAAGACAATTAGCACATGGATCAGCGACGGCCATGTAAAAGA<br>TTTCTATTTATATCCTGGAAAATACACATTCGTCGAGACTGCCGCCCCAGACGGTTAT<br>GAGGTAGCGACTCCGATTGAGTTTACCGTTAATGAAGACGGACAAGTAACCGTTGAT<br>GGAGAAGCCACCGAGGGCGATGCCCACACGTAG |

Red: TEE sequence, Orange: Spy tag, Blue: $His_6$ tag, Green: SUMO tag, Brown: Avi tag, Purple: Protein

**Table S2. Amino acid sequence used for AlphaFold3 prediction**

| chain | Sequence used for the prediction |
|---|---|
| SfbI-TED | DEKTVPNFKSPDPDYPWYGYDSYRGIFARYHNLKVNLKGSKEYQAYCFNLTKYFPRPTYST<br>TNNFYKKIDGSGSAFKSYAANPRVLDENLDKLEKNILNVIYNGYKSNANGFMNGIEDLNAILVT<br>QNAIWYYSDSAPLNDVNKMWEREVRNGEISESQVTLMREALKKLIDPNLEATAANKIPSGYR<br>LNIFKSENEDYQNLLSAEYVP |
| Fg-Aα | ACKDSDWPFCSDEDWNYKCPSGCRMKGLIDEVNQDFTNRINKLKNSLFEYQKNNKDSHSL<br>TTNIMEILRGDFSSANNRDNTYNRVSEDLRSRIEVLKRKVIEKVQHIQLLQKNVRAQLVDMKR<br>LEVDIDIKIRSCRGSCSRALAREVDLKDYEDQQKQLEQVIAKDLLPSRDRQ |
| Fg-Bβ | KAPDAGGCLHADPDLGVLCPTGCQLQEALLQQERPIRNSVDELNNNVEAVSQTSSSSFQY<br>MYLLKDLWQKRQKQVKDNENVVNEYSSELEKHQLYIDETVNSNIPTNLRVLRSILENLRSKIQ<br>KLESDVSAQMEYCRTPCTVS |
| Fg-γ | RFGSYCPTTCGIADFLSTYQTKVDKDLQSLEDILHQVENKTSEVKQLIKAIQLTYNPDESSKP<br>NMIDAATLKSRKMLEEIMKYEASILTHDSSIRYLQEIYNSNNQKIVNLKEKVAQLEAQCQEPCK<br>DTV |

**Table S3. DNA Primer sequences for FASTIA**

| Name | Sequence |
|---|---|
| D83N_F | GGTACGGGTACAATTCCTATCGTGGTATTTTTGCACG |
| D83N_R | GAATTGTACCCGTACCATGGATAGTCAGGATCTGG |
| S84A_F | GGTACGGGTACGATGCCTATCGTGGTATTTTTGCAC |
| S84A_R | GGCATCGTACCCGTACCATGGATAGTCAGGATC |
| Y85A_F | GGTACGGGTACGATTCCGCACGTGGTATTTTTGCAC |
| Y85A_R | GGAATCGTACCCGTACCATGGATAGTCAGGATC |
| R86A_F | CGGGTACGATTCCTATGCAGGTATTTTTGCACGCTACC |
| R86A_R | TAGGAATCGTACCCGTACCATGGATAGTCAGGATC |
| G87A_F | CGGGTACGATTCCTATCGTGCAATTTTTGCACGCTAC |
| G87A_R | GATAGGAATCGTACCCGTACCATGGATAGTCAGG |
| I88A_F | CCTATCGTGGTGCATTTGCACGCTACCATAATCTGAAGG |
| I88A_R | ATGCACCACGATAGGAATCGTACCCGTACCATGGATAG |
| F89A_F | CCTATCGTGGTATTGCAGCACGCTACCATAATCTGAAGG |
| F89A_R | CAATACCACGATAGGAATCGTACCCGTACCATGGATAG |
| A90W_F | GGTACGATTCCTATCGTGGTATTTTTTGGCGCTACCATAATCTG |
| A90W_R | GATAGGAATCGTACCCGTACCATGGATAGTCAGG |
| R91A_F | CGATTCCTATCGTGGTATTTTTGCAGCCTACCATAATCTGAAGG |
| R91A_R | CCACGATAGGAATCGTACCCGTACCATGGATAG |
| Y92A_F | GCACGCGCCCATAATCTGAAGGTTAATCTTAAAGGTTCGAAG |
| Y92A_R | ATTATGGGCGCGTGCAAAAATACCACGATAGGAATCGTACC |
| H93A_F | GCACGCTACGCAAATCTGAAGGTTAATCTTAAAGGTTCGAAGG |
| H93A_R | GATTTGCGTAGCGTGCAAAAATACCACGATAGGAATCGTACC |
| H93F_F | GCACGCTACTTTAATCTGAAGGTTAATCTTAAAGGTTCGAAGG |
| H93F_R | GATTAAAGTAGCGTGCAAAAATACCACGATAGGAATCGTACC |
| L112A_F | GCAACAAAGTACTTCCCGCGTCCCACTTACTCAACC |
| L112A_R | GGAAGTACTTTGTTGCATTGAAACAATACGCCTGGTATTCCTTCG |
| T113A_F | GCAAAGTACTTCCCGCGTCCCACTTACTCAACC |
| T113A_R | CGCGGGAAGTACTTTGCTAAATTGAAACAATACGCCTGG |
| K114A_F | GCATACTTCCCGCGTCCCACTTACTCAACCACG |
| K114A_R | GACGCGGGAAGTATGCTGTTAAATTGAAACAATACGCCTGG |
| Y115A_F | GGCCTTCCCGCGTCCCACTTACTCAACCAC |
| Y115A_R | GGACGCGGGAAGGCCTTTGTTAAATTGAAACAATACGCCTG |
| F116A_F | CGCCCCGCGTCCCACTTACTCAACCACG |
| F116A_R | GTGGGACGCGGGGCGTACTTTGTTAAATTGAAACAATACG |
| P117A_F | GTACTTCGCACGTCCCACTTACTCAACCACGAAC |
| P117A_R | GGACGTGCGAAGTACTTTGTTAAATTGAAACAATACGCCTGG |
| R118A_F | GTACTTCCCGGCACCCACTTACTCAACCACGAAC |
| R118A_R | GGTGCCGGGAAGTACTTTGTTAAATTGAAACAATACGCCTGG |
| P119A_F | GTACTTCCCGCGTGCCACTTACTCAACCACG |
| P119A_R | GCACGCGGGAAGTACTTTGTTAAATTGAAACAATACGCCTG |
| T120A_F | CCCGCGTCCCGCATACTCAACCACGAAC |
| T120A_R | TATGCGGGACGCGGGAAGTACTTTGTTAAATTGAAACAATACG |
| Y121A_F | CCGCGTCCCACTGCCTCAACCACGAACAAC |
| Y121A_R | GAGGCAGTGGGACGCGGGAAGTACTTTGTTAAATTGAAAC |
| S122A_F | CGCGTCCCACTTACGCAACCACGAACAACTTC |
| S122A_R | GCGTAAGTGGGACGCGGGAAGTACTTTGTTAAATTGAAAC |
| T123A_F | GCGTCCCACTTACTCAGCCACGAACAACTTCTAC |
| T123A_R | GAGTAAGTGGGACGCGGGAAGTACTTTGTTAAATTGAAAC |

**Table S3 (continued)**

| Name | Sequence |
|---|---|
| T124A_F | GCAAACAACTTCTACAAAAAGATTGACGGGTCCGGCAG |
| T124A_R | GTAGAAGTTGTTTGCGGTTGAGTAAGTGGGACGCGGGAAG |
| N125A_F | GGCCAACTTCTACAAAAAGATTGACGGGTCCGGCAG |
| N125A_R | TTGTAGAAGTTGGCCGTGGTTGAGTAAGTGGGACGCGGGAAG |
| P199A_F | GGCCTTGAACGATGTAAATAAAATGTGGGAGCGCGAG |
| P199A_R | ACATCGTTCAAGGCCGCTGAATCTGAGTAATACCAAATCG |
| L200A_F | GCCCGCAAACGATGTAAATAAAATGTGGGAGCGCGAG |
| L200A_R | ACATCGTTTGCGGGCGCTGAATCTGAGTAATACCAAATCG |
| N201A_F | GGCCGATGTAAATAAAATGTGGGAGCGCGAGGTCC |
| N201A_R | TTATTTACATCGGCCAAGGGCGCTGAATCTGAGTAATACCAAATC |
| D202A_F | GAACGCAGTAAATAAAATGTGGGAGCGCGAGGTCC |
| D202A_R | TTATTTACTGCGTTCAAGGGCGCTGAATCTGAGTAATACCAAATC |
| V203A_F | GAACGATGCAAATAAAATGTGGGAGCGCGAGGTCC |
| V203A_R | TTATTTGCATCGTTCAAGGGCGCTGAATCTGAGTAATACCAAATC |
| N204A_F | GCAAAAATGTGGGAGCGCGAGGTCCGTAATGGTG |
| N204A_R | GCTCCCACATTTTTGCTACATCGTTCAAGGGCGCTG |
| K205A_F | GCAATGTGGGAGCGCGAGGTCCGTAATGGTG |
| K205A_R | GCGCTCCCACATTGCATTTACATCGTTCAAGGGCGCTG |
| M206A_F | GCATGGGAGCGCGAGGTCCGTAATGGTGAG |
| M206A_R | CTCGCGCTCCCATGCTTTATTTACATCGTTCAAGGGCGCTG |
| W207A_F | GGCAGAGCGCGAGGTCCGTAATGGTGAGATC |
| W207A_R | GGACCTCGCGCTCTGCCATTTTATTTACATCGTTCAAGG |
| E208A_F | GTGGGCACGCGAGGTCCGTAATGGTGAG |
| E208A_R | GACCTCGCGTGCCCACATTTTATTTACATCGTTCAAGG |
| R209A_F | GTGGGAGGCCGAGGTCCGTAATGGTGAGATC |
| R209A_R | GGACCTCGGCCTCCCACATTTTATTTACATCGTTCAAGG |
| E210A_F | GTGGGAGCGCGCAGTCCGTAATGGTGAG |
| E210A_R | GACTGCGCGCTCCCACATTTTATTTACATCGTTCAAGG |
| PUREfrex-F | GAAATTAATACGACTCACTATAGGGAGATCACAACG |
| PUREfrex-R | GGCTTTGTTAGCAGCCGGATCTC |
